# Mean yield gap explains differences in yield stability between organic and conventional systems

**DOI:** 10.64898/2026.06.04.730123

**Authors:** Tamara Ben-Ari, David Makowski

## Abstract

Over recent decades, numerous studies have compared agricultural productivity across different management systems and environments, with increasing attention to the stability of crop yields. Yield stability is a key component of agroecosystem resilience under global change, yet it remains unclear whether some cropping systems are more stable than others. The debate over organic farming is particularly heated, as some evidence indicates that organic systems are less stable, while other evidence suggests the opposite.

Here, we develop a conceptual and statistical framework for comparing crop yield stability across agricultural systems that explicitly accounts for a parametric function relating inter-annual yield standard deviation to mean yield value. Using a new global dataset of long-term field experiments spanning multiple crop species and environments, we show that the fitted functions relating yield standard deviation and mean yield value obtained for organic and conventional cropping systems share several important similarities. For both systems, the yield coefficient of variation (CV) significantly decreases with mean yield, and most of the difference in yield stability between conventional and organic systems is explained by differences in mean productivity. As a consequence, the CV of the two systems becomes similar as the difference in average efficiency narrows.

Our framework reconciles apparently conflicting results in the literature and provides a more robust basis for comparing stability across cropping systems.

## 1 Introduction

Comparing the interannual stability of crop yields across agricultural systems remains both conceptually and statistically challenging. As formalized by Taylor’s power law (TPL) [1], and widely documented across biological and non-biological systems [2, 3, 4, 5], including crop yields [6], the variance of a quantity typically scales with its mean. Thus, changes in average crop yield are intrinsically associated with changes in yield variance, implying that apparent differences in stability may simply reflect differences in mean productivity rather than genuine differences in underlying variability.

Stability is often quantified using the coefficient of variation (CV), defined as the ratio of the standard deviation to the mean. When the standard deviation is proportional to the mean, the CV is constant and independent of system average productivity. However, when this proportionality does not hold, CV-based comparisons may be misleading, as differences in CV can depend on underlying differences in mean yield. Consequently, when two cropping systems differ in their mean yields and when their CV depend on these values, differences in CV may simply reflect differences in average productivity rather than intrinsic differences in variability. In such cases, comparisons of yield stability must explicitly account for the relationship between CV and mean yield.

The question of inter-annual yield stability is particularly salient in the long-standing debate over the performance of organic versus conventional agriculture. While recent syntheses consistently report a persistent organic-to-conventional average yield loss (or yield gap), typically ranging from approximately −16% to −34%, depending on crop species, management practices, and environmental context [7, 8, 9]. An increasing number of studies have examined whether organic and conventional systems differ in their temporal and spatial yield stability, both in local experiments (see for example, [10, 11, 12, 13, 14]) and in global syntheses [15, 16, 17]. These comparisons are often motivated by the assumption that organic and conventional management practices generate inherent differences in stability due to contrasting levels of regulation by biological processes and differences in crop sensitivities to pests,diseases, and nutrient deficits.

From a theoretical perspective, arguments support both higher and lower yield stability in organic systems relative to conventional systems. On the one hand, organic agriculture promotes greater biological and functional diversity at field and landscape scales, which, according to ecological theory and empirical evidence from natural ecosystems, can enhance stability through insurance and complementarity effects [18, 19]. Empirical studies of diversified farming systems also suggest that these stabilizing mechanisms are at play [20, 21]. Organic management is consistently associated with improvements in soil properties, including higher soil organic matter content [22], increased biological activity [23, 24], and improved physical structure [25]. Such changes can buffer crops against climatic extremes by enhancing water-holding capacity, nutrient retention, and root penetration, thereby contributing to greater resilience under drought or heavy rainfall events. On the other hand, organic systems may exhibit lower yield stability because of the negative impacts of weeds, pests, and diseases, potentially leading to more frequent or severe yield losses [26]. Nutrient management may also be less effective in organic systems, as nutrient availability from organic inputs is highly variable, whereas mineral fertilizers in conventional systems provide a more immediate and predictable nutrient supply [27, 28], potentially translating into lower stability of yields over time and space.

Various metrics are commonly used to study the stability of different ecosystems, particularly variance, standard deviation, and the coefficient of variation (CV). Several metaanalyzes have compared yield stability between organic and conventional systems across large collections of experiments. Lesur-Dumoulin et al. (2017) found no significant difference in temporal or spatial stability, quantified using yield variances, for a dataset of horticultural crop yields [16]. Knapp and van der Heijden, reanalyzing a global dataset of cereal trials, similarly reported no difference in variances but concluded that relative stability, measured by yield CV, was on average higher (lower CV) in conventional systems by approximately 15% (range: 2–30%) [15]. Smith et al. (2019) extended this approach to include associated biodiversity and likewise reported higher relative stability (lower CV) in conventional systems [17].

Concerning crop yield stability, the CV is probably the most popular metric for comparing different cropping systems, especially in the context of agroecology and agricultural sustainability [29, 30, 31]. The popularity of the CV stems mainly from the fact that this metric is a measure of yield variability, expressed relative to an average yield, enabling cross-crop comparisons. For this reason, the CV is often considered independent of the average yield, although this is not necessarily the case. In fact, in the next section, we show that although the CV remains independent of the mean yield when the standard deviation is strictly proportional to the mean (on a linear scale), this property no longer holds when strict proportionality does not apply. We analytically demonstrate that when the standard deviation is not strictly proportional to the mean, the value of the CV is no longer independent of the mean yield value. Then, CV-based comparisons cannot disentangle intrinsic differences in variability between systems from the mechanical effect of mean yield differences. In this case, differences in CV may reflect genuine stability contrasts between management systems or be driven by underlying mean yield differences. These considerations are particularly important to keep in mind when comparing organic and conventional systems because average yields have been shown to be consistently lower in organic farming. Such differences in average yields must be taken into account when interpreting differences in CV between organic and conventional systems.

Here, we strive to analytically and empirically disentangle yield variability from mean productivity. Specifically, we (i) develop an analytical framework to assess yield stability in contrasted agricultural systems while accounting for the relationship between CV and mean yield; (ii) apply this framework to quantify and compare the relationships between mean yields and yield CV in organic and conventional systems using a new global dataset of mid-to long-term experiments spanning multiple crop species, locations, and management contexts (minimum duration *≥* four years per crop–site combination); and (iii) evaluate the contribution of key agronomic and environmental factors to differences in stability between organic and conventional systems. Our approach provides a robust basis for disentangling intrinsic stability properties from differences driven by average yield levels and offers a more general interpretation of previous empirical findings.

## 2 A framework for comparing yield stability based on Taylor’s power law

### 2.1 Taylor’s power law

In many biological, physical, and ecological processes, the variance or standard deviation of a quantity scales with its mean according to Taylor’s power law (TPL). Here, we use this law to relate yield standard deviation to mean yield as follows:

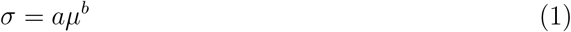

where *µ* is the inter-annual mean or expected yield, *σ* is the yield standard deviation (measuring between-year yield variability) in spatial unit *i*, and *a* and *b* are parameters to be estimated (see methods).

Taking logarithms, we obtain a linear relationship:

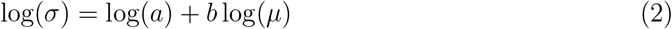

When yield time series are available for organic and conventional cropping systems, a separate Taylor’s power law (TPL) can be fitted by linear regression (see methods), leading to a parametric *σ* -*µ* relationship for organic systems, log(*σ*_org_) = log(*a*_org_) + *b*_org_ log(*µ*_org_), and another one for conventional systems, log(*σ*_conv_) = log(*a*_conv_) + *b*_conv_ log(*µ*_conv_).

Depending on the values of Taylor’s power law parameters, four possible patterns can be defined regarding the relative position of the *σ* -*µ* relationships of the two systems, which are illustrated schematically in Figure 1. First, both systems may share the same scaling relationship (*i*.*e*., *a*_org_ = *a*_conv_ = *a* and *b*_org_ = *b*_conv_ = *b*; case (a)). In this situation, both systems have equal standard deviation *σ* if they have equal mean yield, and any difference in standard deviation arises solely from differences in mean yield, such that lower-yielding systems mechanically exhibit lower standard deviation *σ* and thus appear more stable according to this metric. Second, systems may share the same slope but differ in intercept (*i*.*e*., *b*_org_ = *b*_conv_ and *a*_org_ ≠ *a*_conv_; cases (b)–(c)). In this case, one system exhibits systematically higher or lower standard deviation *σ* across the entire range of mean yield values, and the system with the lower intercept is more stable according to *σ* for any given mean yield. Finally, both the slope and the intercept may differ between systems (*i*.*e*., *a*_org_≠ *a*_conv_ and *b*_org_≠ *b*_conv_; case (d)). Under this configuration, the difference of *σ* between the two systems depends on their relative mean yields, and the ranking of the systems according to *σ* may even reverse across the mean yield gradient. Determining which of these configurations best describes the relationship between mean yield and standard deviation is, therefore, essential for accurately characterizing the relative stability of organic and conventional systems.

**Figure 1:**
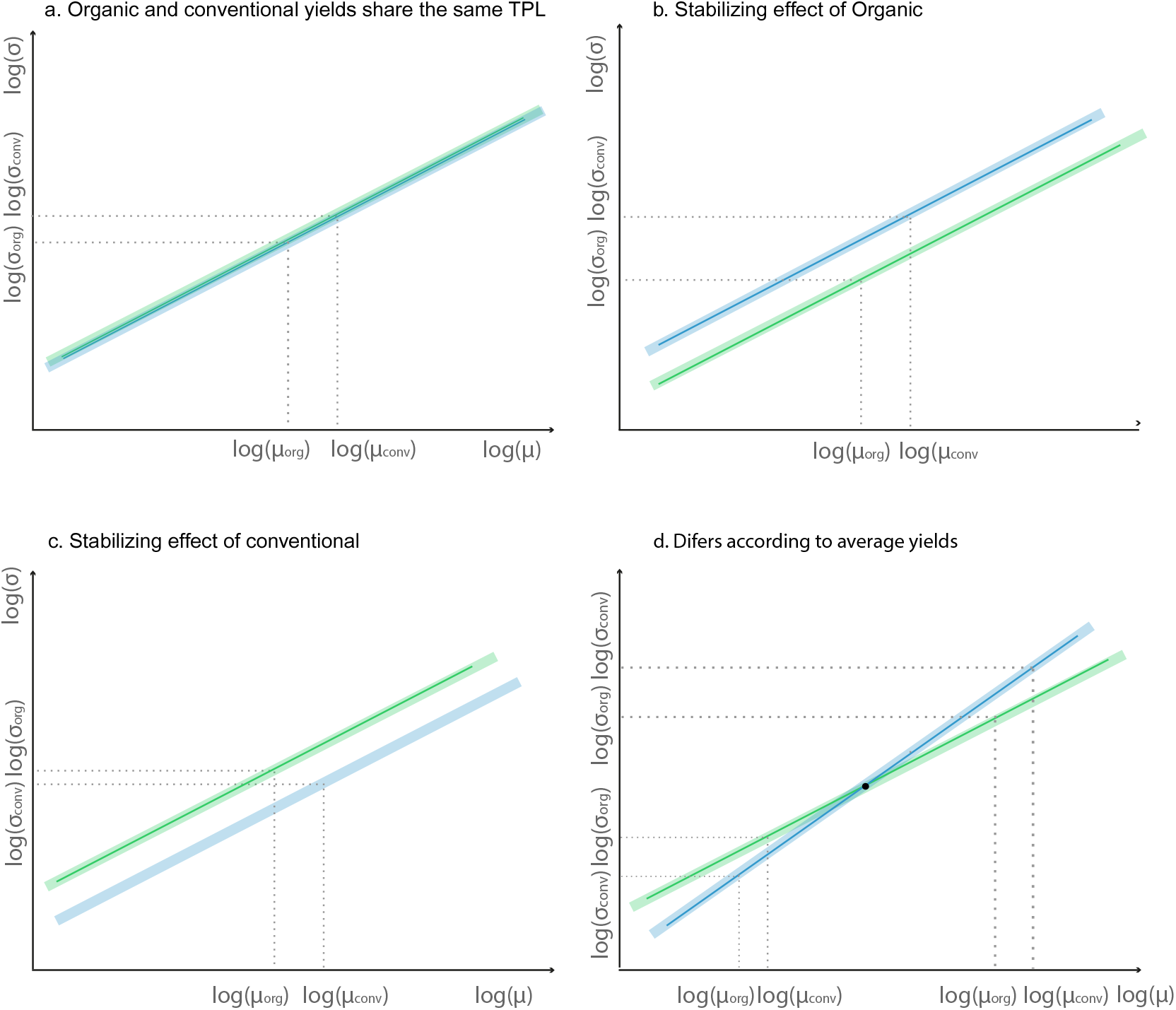
Schematic representation of four different configurations of organic and conventional Taylor’s power laws (TPLs). TPL describes the relationship between mean yield (*µ*) and yield standard deviation (*σ*; Eq. 1). In its log-transformed form, this relationship is assumed to be linear (Eq. 2). The green line and shaded area represent an hypothetical TPL for organic systems, whereas the blue line and shaded area represent a hypothetical TPL for conventional systems. gray dotted lines indicate, schematically, the mean yield and standard deviation assuming *µ*_org_ *< µ*_conv_. This framework leads to four distinct configurations: (i) the two systems may share the same TPL (panel a), (ii) differ only in intercept (panels b and c), or (iii) differ in both intercept and slope (panel d). A symmetric counterpart to panel (d), in which the organic slope is steeper than the conventional slope, is also possible but is not shown here. Each configuration has different implications for relative stability. In panels (a) and (b), organic yield standard deviation is expected to be lower because organic mean yield is lower. In panel (c), organic standard deviation may be higher despite lower mean yields. In panel (d), the relative stability of the two systems depends on the magnitude of the yield difference between the two systems and on the difference between scaling parameters.

### 2.2 Why CV-based comparisons are sometimes misleading

The coefficient of variation (CV) is often used as a measure of relative yield stability. It is thus important to understand the implications of the TPL for the interpretation of differences in CV between cropping systems. Under the Taylor power Law, the coefficient of variation is expressed as:

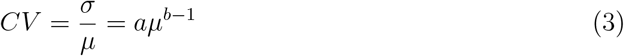

This expression shows that the yield CV is independent of the mean yield only if *b* = 1 (linear proportionality), in which case the coefficient of variation remains constant across all levels of mean yield, *i*.*e*.,

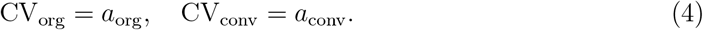

On the contrary, when *b* ≠ 1, the CV is no longer independent of the mean yield. Differences in mean yield alone are sufficient to generate differences in CV, even if two systems share the same scaling relationship (*i*.*e*.,*b*_org_ = *b*_conv_).

This dependence can be made explicit by expressing the ratio of coefficients of variation between systems as:

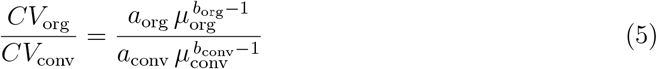

This expression can be rewritten as:

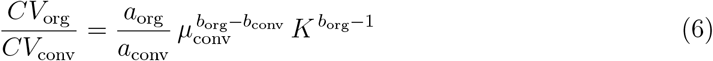

where *K* is the mean yield ratio, defined as:

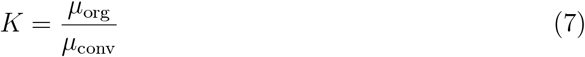

This formulation shows that differences in relative stability between organic and conventional systems (*i*.*e*., differences in CV) depend jointly on the parameters of Taylor’s power law (*a* and *b*), the mean yield ratio between systems *K* (*i*.*e*., relative average yield gap), and, when the two systems have different parameters *b*, the mean yield of the conventional system through *µ*_conv_ (Eq. 6).

To better understand the implications of this relationship, it is useful to consider a set of limiting cases, as presented in Figure 2. First, consider the situation where organic and conventional systems share the same TPL slope (*i*.*e*., *b*_org_ = *b*_conv_ = *b*; see cases (a)–(c) in Figure 1). In this case, equation 6 simplifies to:

**Figure 2:**
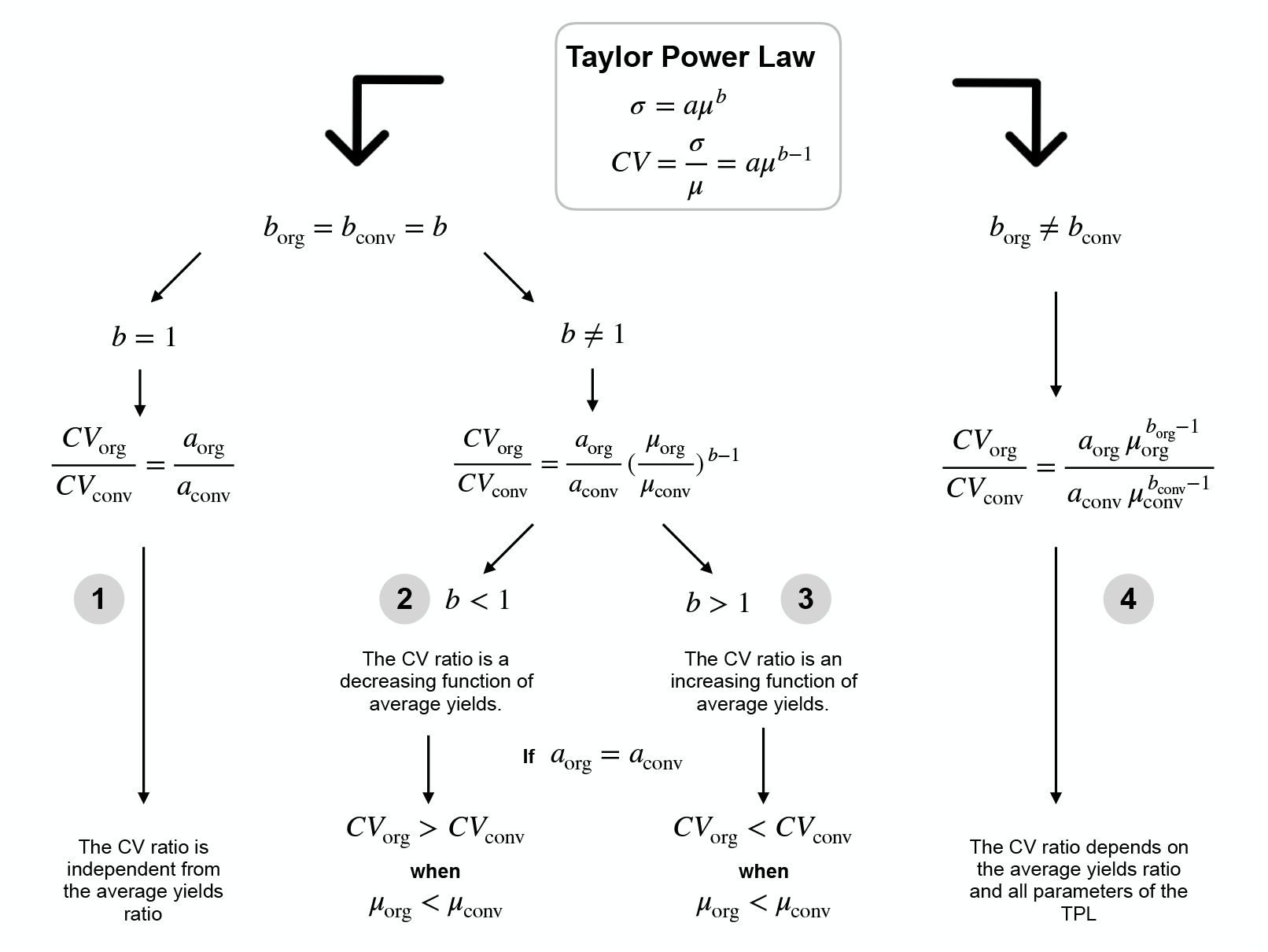
Implications of Taylor’s power law for comparing yield stability based on the coefficient of variation (CV). Under Taylor’s power law (TPL), yield standard deviation scales with mean yield as *σ* = *aµ*^*b*^, implying that the CV depends on mean yield according to *CV* = *aµ*^*b™*1^. When organic and conventional systems share the same TPL slope (*i*.*e*.,*b*_org_ = *b*_conv_ = *b*), the ratio of CV is independent from the mean only if *b* = 1 (case 1). If *b <* 1, the CV is a decreasing function of mean yield, while CV is an increasing function of mean yields if *b >* 1.Finally, when the TPL slopes are different between the two systems (case 4), the ratio of CV depends jointly on four parameters (*i*.*e*., *a*_org_, *a*_conv_, *b*_org_ and *b*_conv_) and on the yield ratio between systems. This framework shows that comparisons of relative stability based on the CV cannot be interpreted independently from the mean yields, unless *b* = 1.

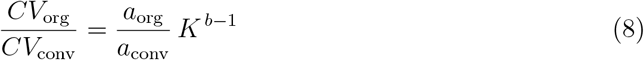

If *b* = 1, the CV becomes independent of the mean yield, and we obtain: *CV*_org_ =*a*_org_, *CV*_conv_ = *a*_conv_ or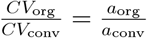. The CV ratio is therefore independent of the ratio of average yields, and the conclusion regarding the relative stability of the two systems remains the same, regardless of the average yields of organic and conventional systems.

If *b* ≠ 1, *CV* depends on mean yield and the ratio of *CV* to the mean yield ratio *K*. If *b <* 1, the CV is a decreasing function of mean yield ratio. Conversely, if *b >* 1, the CV increases with mean yield ratio.

To summarize, when organic and conventional systems share the same TPL parameter *b* (*i*.*e*., the slope of the TPL when expressed as a log), differences in CV are independent of differences in mean yield only if *b* = 1, *i*.*e*., if the standard deviation is proportional to the mean yield (*σ*_*org*_ = *a*_*org*_*µ*_*org*_ and *σ*_*conv*_ = *a*_*conv*_*µ*_*conv*_). When *b* is different from 1 or when organic and conventional systems have different values of *b*, the ratio of the CV of the two systems is no longer independent of mean yields. Any conclusion about the relative stability of one system compared to another may then depend on the mean yield values themselves (see Figure 2). Conclusions based on CV are thus independent from mean yields only in a very specific situation, *i*.*e*., when the slopes of the TPL of the two systems are both equal to 1.

### 2.3 Implications for comparing cropping systems using CV

When CV depends on mean yield and when cropping systems have contrasting mean yield levels, differences in CV between cropping systems should be interpreted with care because such differences may simply reflect differences in mean yield values. The effect of the mean value on CV is a direct implication of the TPL, and it occurs as soon as the yield standard deviations of the systems are not strictly proportional to the mean yields. These considerations highlight the necessity to account for *σ*-*µ* relationships when comparing yield stability between two systems based on CV. In the next section, we propose a TPL model relating *σ* to *µ*, including the type of cropping system as a covariate. The model is fitted to experimental data, and the fitted model is then used to compute the ratio of CV of the two systems as a function of their mean yield ratio. The results are used to analyze the sensitivity of CV ratio to mean yield ratio.

## 3 Materials and Methods

### 3.1 Data

We use a global database of long-term field experiments comparing organic, diversified, and conventional agricultural systems, recently compiled and made openly available by Sawadogo and Ben-Ari [32]. The dataset was constructed by merging five existing meta-analytic databases and conducting a systematic literature review over the time frame 1950-2025. Studies were selected based on strict criteria to ensure comparability: yield measurements for the same crop species, at the same site, under comparable agronomic conditions, and over a minimum duration of four years. The final database includes 102 peer-reviewed studies conducted at 73 experimental sites across 23 countries and six continents, corresponding to 322 paired comparisons between alternative (organic or diversified) and conventional systems. Experimental durations range from 4 to 29 years. The dataset spans 39 crop species grouped into nine crop types, with a predominance of cereals, and includes harmonized metadata describing crop characteristics, geographic context, experiment duration, and key management practices (*i*.*e*., on-farm versus controlled experiment, multi versus mono cropping, rotation, irrigation, tillage, and fertilization levels, see Supplement section S1.1).

Our unit of analysis considered to compute mean yields and inter-annual standard deviations is the experimental unit (EU), defined as a unique combination of site, crop species, and management system. Each EU consists of a time series of annual yields from which mean yields and standard deviations are derived. This definition ensures statistical independence across observations while preserving the within-study structure. The final dataset comprises 322 EUs distributed across multiple studies and geographic regions (See Supplement section S1.2).

### 3.2 Variables and derived metrics

We quantify differences between organic and conventional systems using three effect size metrics: the log response ratio of mean yields, the log variance ratio, and the log coefficient of variation (CV) ratio. They are calculated for each experimental unit separately. To account for differences in sampling precision among experimental units, all effect sizes are associated with analytically derived sampling variances and bias corrections following [33], see supplementary section S2.1 and S2.

We first quantify overall differences in productivity and variability between organic (including diversified) and conventional systems using intercept-only Bayesian mixed-effects models (see Supplementary section S1). Organic systems exhibit a significant yield gap, with mean yields about 15% lower than conventional systems (yield ratio: 0.85 [0.80–0.91]). In contrast, no significant difference is detected in absolute variability, with variance ratios close to unity (1.01 [0.90–1.12]). We find that the coefficient of variation (CV), is significantly higher in organic systems (CV ratio: 1.18 [1.07–1.31]), corresponding to an increase of approximately 18%. Full definitions of these metrics, including their exact formulations and associated uncertainty estimates, are provided in the Supplementary Information Section SS3.

### 3.3 Statistical modeling

We define Bayesian mixed-effects models to implement our TPL framework (see equation 2. The response variable is the logarithm of the yield standard deviation, and the main predictor is the logarithm of the mean yield. The management system (*i*.*e*., organic or diversified versus conventional) is included as a fixed effect, together with its interaction with mean yield, to test whether the scaling relationship differs between systems. To account for additional sources of variability, we include a set of potential covariates (*Z*_*i*_) describing the agronomic or environmental characteristics of the experimental units (*i*.*e*., crop types, on-farm versus controlled experiments, multi versus mono cropping, rotation, irrigation, tillage, and fertilization levels).

The specification of the basis model is:

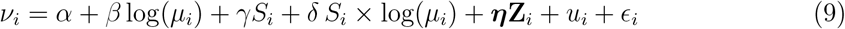

where *v*_*i*_ is the corrected logarithm of the yield standard deviation in the ith EU (eq. (7) of [33]), *µ*_*i*_ is the mean yield, *S*_*i*_ is a dummy variable describing the management system (*S* = 1 for organic, *S* = 0 for conventional), **Z**_*i*_ is a vector of covariates (fixed effects), *u*_*i*_ is a random intercept associated with the experimental unit, and *ϵ*_*i*_ is the residual error term describing the estimation error of the log standard deviation (whose variance is estimated from eq.(8) in [33]).

We compare several variants of the above-model to evaluate whether the relationship is further explained by agronomic or environmental covariates *Z*. We test a broad range of candidate variables available in the dataset, including detailed crop-type classes, a simplified crop-type grouping (“Grain”versus “Garden”), continent, development level, experiment duration (both as a continuous variable and as a categorical duration class), tillage, multicropping, and irrigation. The “Grain” category includes cereals, oil crops, legumes, forage, and fiber crops, whereas the “Garden” category groups fruits, vegetables, and root and tuber crops, based on the fact that these categories are not centered on the same levels of expected yields. For each covariate, we compare models including additive effects only, models allowing the covariate to modify the intercept of the relationship, and models allowing it to interact with mean yield, thereby modifying the slope of the mean–variance relationship.

Models are estimated using the `MCMCglmm` package in R under a Gaussian error distribution. We used weakly informative priors (see section S S1) for fixed and random effects. Markov Chain Monte Carlo chains were run for 100,000 iterations, with a burn-in of 20,000 iterations and a thinning interval of 10. Model comparison is based on the Deviance Infor-mation Criterion (DIC), allowing us to identify the most parsimonious specification among competing models differing in fixed effects and interaction terms, together with the statistical significance of fixed effects. All analyzes were conducted in R (R Core Team, version 4.5.2), using the `MCMCglmm` package for Bayesian mixed-effects modeling.

The final model includes the management system *S* (organic or conventional), mean yield, and their interaction (*S*_*i*_ *×* log(*µ*_*i*_)), allowing the relationship between standard deviation and mean yield to differ between systems, together with a simplified crop-type grouping (‘Grain”versus ‘Garden”), while accounting for variability among experimental units and weighting log standard deviation by their associated variance. For model selection details see supplementary section S1.4.

The selected model is expressed as:

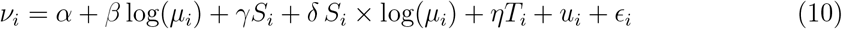

where *v*_*i*_ is the corrected logarithm of the yield standard deviation, *µ*_*i*_ is the mean yield, *S*_*i*_, the management system (organic/diversified vs. conventional), and *T*_*i*_ is a categorical dummy variable distinguishing crop types (*T* = 1 for “Grain”, *T* = 0 for “Garden”).

Model diagnostics indicate good convergence and mixing of the MCMC chains, with low autocorrelation and high effective sample sizes for both fixed and random effects. Standard convergence diagnostics were satisfied, supporting the robustness of parameter estimates (see section S S1).

The parameters of the TPL defined in section 2 (*a*_*conv*_, *a*_*org*_, *b*_*conv*_, *b*_*org*_) are computed for the two crop types separately, from the posterior distributions of the model parameters (*β, γ, δ, η*).

## 4 Results

### 4.1 Differences in mean–variance scaling between agricultural systems

The selected model reveals a strong positive relationship between mean yield and yield standard deviation, consistent with Taylor’s power law (posterior mean of *β* = 0.80 [0.73, 0.87], *p <* 0.001). The main effect of the management system is significant and positive, with organic systems exhibiting higher variability at a given mean yield (*γ* = 0.21, 95% CI: [0.12, 0.31], *p <* 0.001). In addition, the interaction between the management system and mean yield is significant and negative (*δ* = −0.056, 95% CI: [-0.10, -0.008], *p* = 0.018), indicating that the scaling exponent of the TPL differs between systems, with a weaker increase in variability with increasing mean yields in organic systems compared to conventional systems (Fig. 3). This effect is not dominant, as shown by the superimposition of confidence intervals (see supplementary Figure S4). It nevertheless has a small effect on values of *b*_*org*_ and *b*_*conv*_, whose posterior medians are equal to 0.75 and 0.79, respectively (Fig. 3).

**Figure 3:**
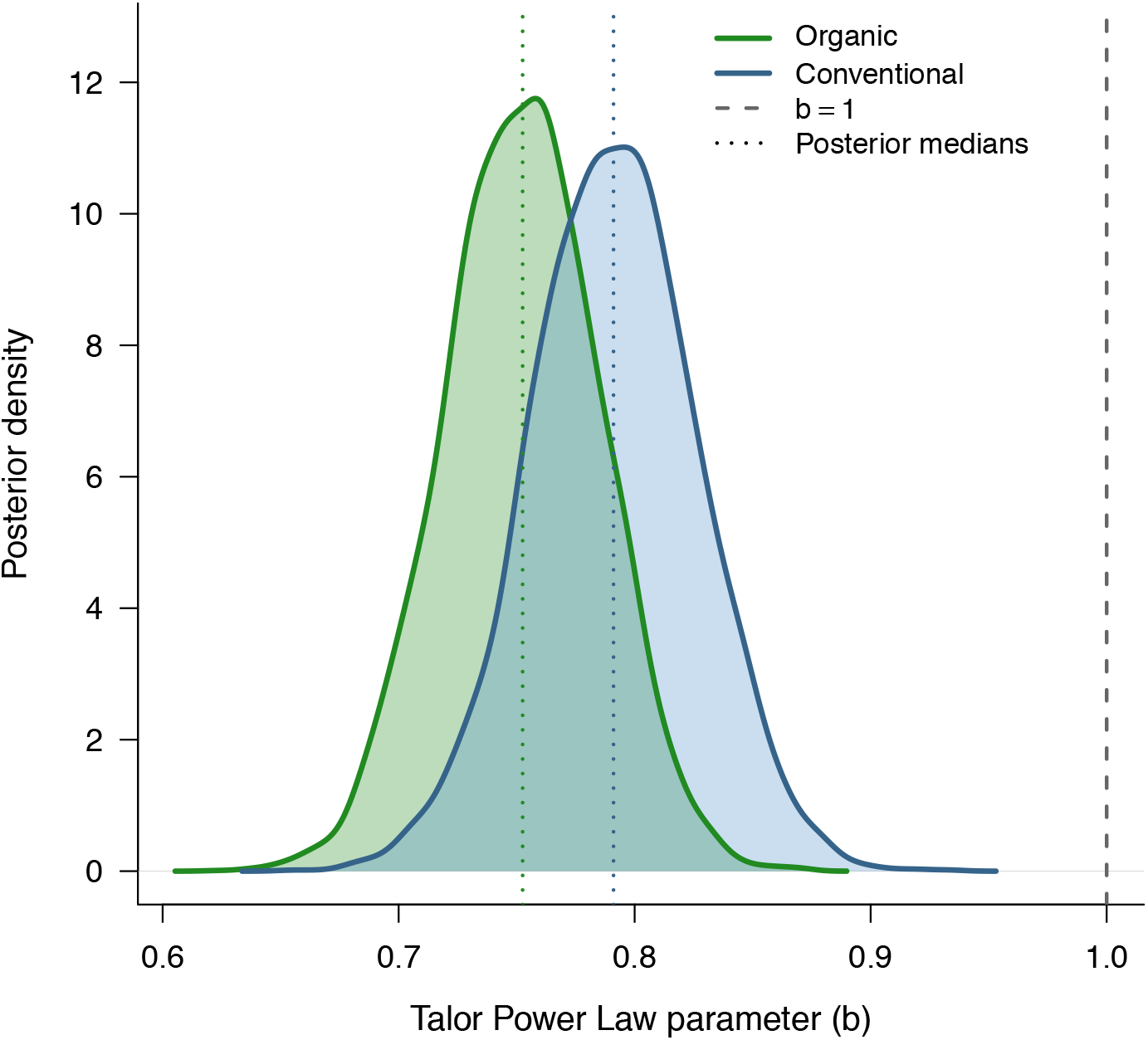
Posterior distributions of the parameter *β* (effect of log mean yield on log standard deviation for conventional system, equal to *b*_*conv*_ in TPL) and of *β* + *δ* (same effect but for organic system, equal to *b*_*org*_ in TPL), generated from 100 000 simulations. The vertical bold gray line indicates the value of 1, corresponding to a proportionality between standard deviation and mean yield. Solid lines represent posterior densities, and thin dashed lines indicating posterior medians (0.75 for organic and 0.79 for conventional systems). Conventional systems exhibit higher values than organic systems, indicating a steeper increase in standard deviation with mean yield.

Grain systems exhibit lower variability than garden systems for a given mean yield (posterior mean of *η* = −0.54, 95% CI: [-0.74, -0.33], *p <* 0.001), highlighting systematic differences across crop groups.

The fitted log–log relationship between yield standard deviation and mean yield is compared across the two cropping systems and the two types of crops in Fig. 4. Consistent with the parameter estimates, conventional systems exhibit a slightly steeper increase in variability with mean yield, whereas organic systems follow a more moderate scaling. Differences between crop groups are reflected in vertical shifts of the relationship, with garden crops generally exhibiting higher variability than grain crops for a given mean yield.

**Figure 4:**
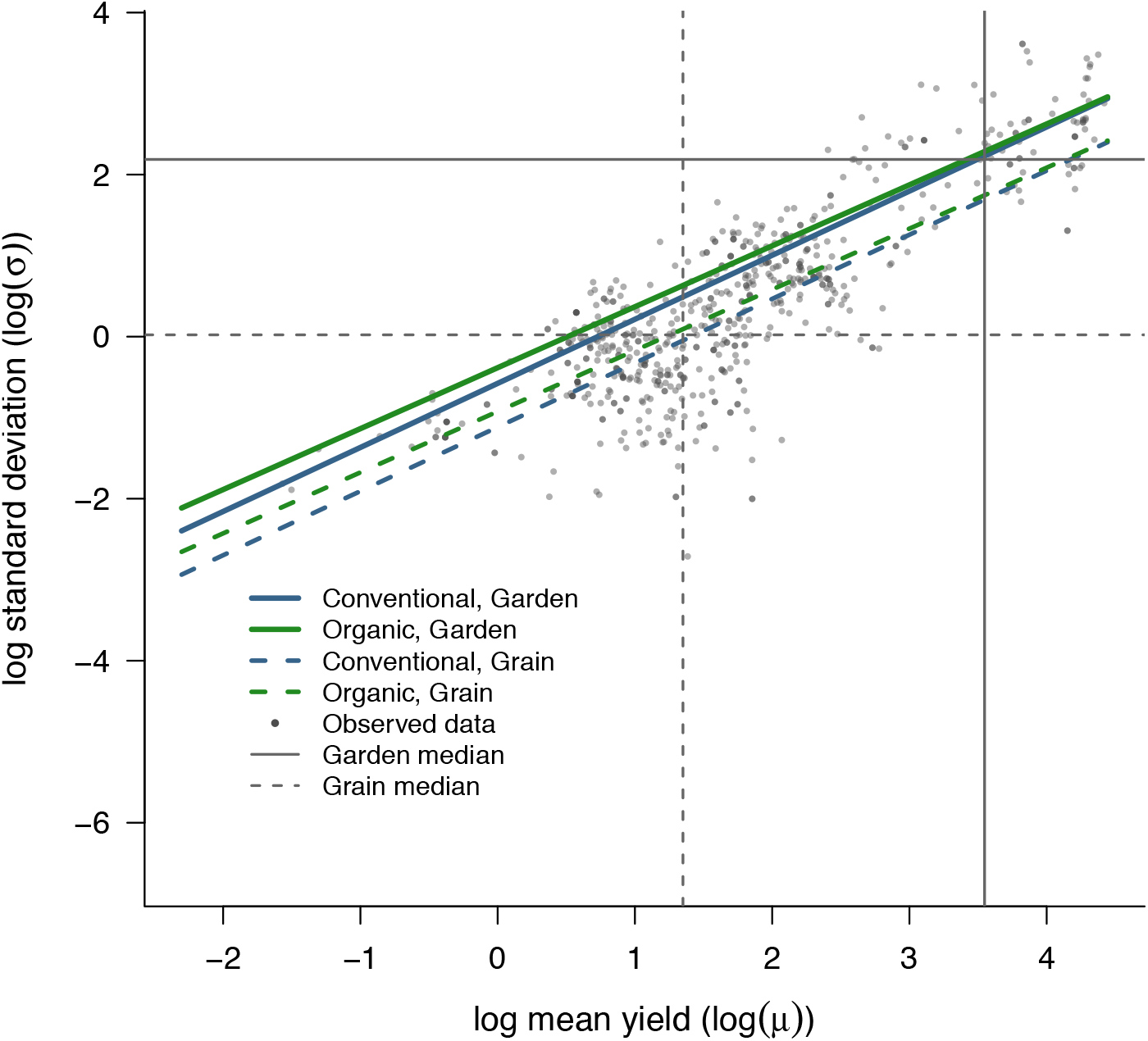
Observed and fitted relationships between mean yield and yield standard deviation on a log–log scale. Points represent experimental units, and lines correspond to fitted relationships for organic and conventional systems across grain and garden crops. Solid lines denote garden crops and dashed lines denote grain crops. Vertical and horizontal lines indicate median values of empirical mean yield and yield standard deviation for each crop group. The figure shows a strong positive relationship between mean yield and yield standard deviation, with conventional systems exhibiting slightly steeper scaling than organic systems and separate intercepts both for organic versus conventional and for grain and garden crop categories.

These results indicate that both the level and the scaling of yield standard deviation differ across agricultural systems, although the magnitude of slope differences remains modest compared to the dominant effect of mean yield. Overall, this figure highlights that both the level and the scaling of standard deviation depend on mean yield.

### 4.2 Dependence of relative stability on yield differences

Building on the mean–standard deviation relationships described above, we next examine how differences in relative stability, measured by the coefficient of variation (CV), depend on differences in mean yield between systems. Because scaling exponents (*b*) are lower than one, the CV decreases with increasing mean yield (see 2 and S5).

Following equation 6, figure 5 illustrates the relationship between CV ratios (organic to conventional) and yield ratios. In both crop groups, the CV ratio decreases as the yield ratio increases, indicating that apparent differences in stability organic vs. conventional diminish as the mean yield gap narrows. Conversely, when organic mean yields are substantially lower than conventional mean yields, organic systems appear less stable, as reflected by higher CV values. Although this pattern is consistent across crop groups, the strength of the relationship differs slightly between grain and garden systems, reflecting differences in the underlying mean-standard deviation relation. Overall, these results demonstrate that CV ratios are intrinsically linked to mean yield differences and therefore cannot be interpreted independently of productivity contrasts between systems.

**Figure 5:**
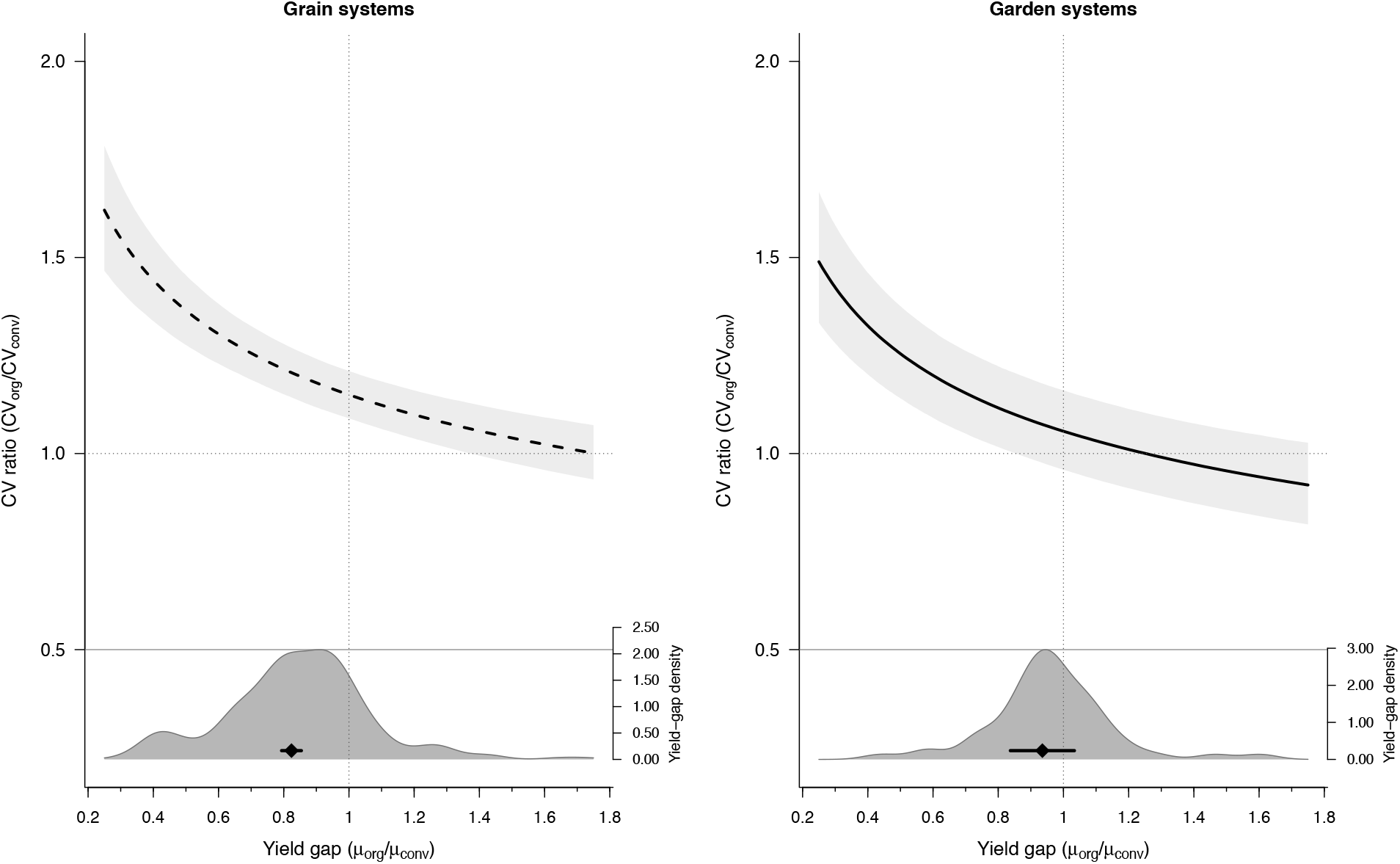
Estimated relation between CV ratio mean yield gap (organic to conventiona) for grain and garden crop groups. Each panel shows the CV ratio, *CV*_org_*/CV*_conv_, as a function of the mean yield ratio, *µ*_org_*/µ*_conv_. The black curve (dashed for “grain” systems and solid for”garden systems” represents the median estimated relationship obtained from the final Taylor power law framework, and the light gray shaded band shows the associated estimated 95% uncertainty interval. The darker gray shaded area at the bottom of each panel represents the empirical probability density of the observed mean yield ratio distribution in the corresponding crop group. The horizontal segment with a diamond indicates the mean yield ratio for that crop group, estimated from an intercept-only Bayesian mixed-effects model fitted to log(*µ*_org_*/µ*_conv_) with experimental units included as a random effect. Confidence intervals are computed taking into account observation-specific measurement-error variances. For “grain” systems, the estimated mean yield ratio was 0.833, with 95% CI [0.769-0.897]. For “garden” systems, the estimated meanyield ratio was 0.923, 95% CI [0.799-1.047]. The vertical dashed line indicates equality in mean yield between systems, *µ*_org_*/µ*_conv_ = 1, and the horizontal dashed line indicates equality in relative variability, *CV*_org_*/CV*_conv_ = 1.

### 4.3 Effects of covariates

To assess the potential influence of additional explanatory variables on yield variability, we extended the selected Taylor’s Power Law (TPL) model by introducing covariates one at a time. Starting from the best-supported model (including the interaction between the management system and the TPL slope, and a main effect of crop type), each covariate was added individually as an additional fixed effect, and model performance was compared using the Deviance Information Criterion (DIC). For each covariate, models were fitted both with and without the interaction between the management system and the slope of the TPL, and the best-performing specification was retained for interpretation.

Among the tested covariates, only a limited subset showed consistent and statistically supported effects on variability. Trial duration had a small but significant positive effect on log(*σ*) (posterior mean = 0.016, 95% CI [0.004; 0.029], *p* = 0.0098), indicating that longer experiments tend to exhibit higher standard deviation at a given mean yield. This effect may reflect the fact that longer trials have a greater probability of integrating climatic extremes. In the dataset, trial duration was included as a continuous covariate (number of years), derived from the reported duration of each experimental comparison. Trial duration was also tested as a categorical variable (*≤* 5, 6⌣10, *>* 10 years), yielding consistent patterns (results not shown)).

Irrigation was associated with a reduction in variability (posterior mean = −0.157, 95% CI [−0.305; 0.003], *p* = 0.0485), suggesting a modest stabilizing effect of water availability. A strong regional effect was observed for Australia, where variability was substantially higher than in the reference region (posterior mean = 0.946, 95% CI [0.429; 1.428], *p* = 0.0005).

All coefficients are expressed on the log scale of the response variable (i.e., log(*σ*)), such that positive values indicate an increase in variability and negative values a decrease, conditional on mean yield and other covariates in the model. In contrast, no robust effects were detected for legume status, level of development, or crop rotation. Overall, these results indicate that variability is primarily shaped by specific environmental and experimental conditions, rather than by broad categorical factors across systems.

## 5 Discussion

The question of yield stability is part of a broader debate on the sustainability and resilience of future cropping systems. In particular, a central issue is whether agricultural diversification and the reliance on sustainable agronomic practices enhance production stability, or whether a higher reliance on inputs (especially mineral fertilizers) in intensified systems leads to reduced yield variability. However, despite decades of empirical research, the comparative stability of organic and diversified versus conventional systems remains debated. In this context, clarifying the drivers of yield stability differences is essential to reconcile conflicting findings.

A central result of our study is that apparent differences in yield stability between organic and conventional systems are primarily driven by differences in mean productivity. Once the dependence between variability and mean yield is explicitly accounted for, evidence for intrinsic differences in variability between systems becomes much weaker. Our results can be directly compared to those of [15], who reported lower relative stability in organic systems based on CV ratios. We recover a similar qualitative pattern, but show that this cannot be interpreted as evidence of intrinsically lower stability. We argue that higher CV values in organic systems primarily reflect lower mean yields. This interpretation is consistent with the absence of differences in absolute variability reported in previous studies [15, 16]. Overall, our results show that reported differences in stability between organic and conventional systems are consistent with similar mean–standard deviation relations across systems, with only modest differences in scaling parameters. The estimated scaling exponents differ slightly between organic and conventional systems but remain close in magnitude, indicating that agronomic practices primarily affect productivity levels rather than fundamentally altering the relationship between mean yield and standard deviation. Taken together, these results indicate that differences in stability largely emerge from differences in mean yield rather than intrinsic variability dynamics.

This result provides a direct explanation for previously reported differences in CV. As a consequence, differences in CV values largely emerge from differences in mean yield. In particular, the CV ratio decreases as the organic-to-conventional mean yield gap narrows, implying that improving organic yields can simultaneously enhance yield stability. This highlights that closing the yield gap in organic systems may constitute a “win–win” outcome, improving both productivity and yield stability.

More generally, these findings can be interpreted within a broader theoretical framework linking productivity and variability. Our results suggest that the trade-off between productivity and stability is governed by a common underlying scaling relationship rather than fundamentally different stability properties across systems. This finding supports the idea that productivity and stability are jointly constrained by shared biophysical processes. While cropping systems and crop types may influence the position along this trade-off, they appear to alter its overall shape only marginally.

While our results highlight a dominant statistical relationship, understanding the underlying mechanisms remains essential. The mechanisms underlying the productivity–stability relationship are likely to be complex and context-dependent. Beyond field-level processes, landscape composition and configuration can influence crop performance through pest regulation, pollination, and microclimatic buffering. While most studies have focused on mean yields, these landscape effects may also influence yield variability by stabilizing or destabilizing ecological processes over time [34]. For instance, agricultural intensification through monocropping may increase vulnerability to environmental stresses, for example, by reducing structural diversity or buffering capacity. Conversely, agroecological principles emphasize diversification, soil organic matter management, and landscape complexity as key drivers of resilience [35]. Similarly, high-yielding varieties optimized for intensive systems may exhibit increased sensitivity to climatic stress [36]. These mechanisms suggest that maximizing productivity and stability can be competing objectives, although the shape of this trade-off may be modulated by management practices.

However, these interpretations must be considered in light of important limitations in the available data. One reason for the lack of convergence in the literature lies in the large uncertainties associated with empirical comparisons. As highlighted in our analysis, available datasets remain limited in size and representativeness. Ideally, comparisons between organic and conventional systems should be conducted under strictly controlled factorial experimental designs, repeated over time and across a wide range of environmental conditions. In addition, experimental plots should be sufficiently large to capture key ecological processes (*e*.*g*., soil biodiversity, landscape interactions) and avoid artifacts related to plot proximity or management spillovers. Importantly, several studies have shown that yield gaps and variability patterns observed at the plot scale may not translate directly to field-scale conditions, where greater environmental heterogeneity and management complexity can amplify both mean differences and variability [37]. This suggests that plot-based experiments may bias real-world variability and bias stability comparisons. Finally, because yield variability is inherently more difficult to estimate than mean yields, long-term experiments are essential to robustly assess stability. Meeting all these requirements entails substantial logistical and financial constraints, which partly explains the persistence of uncertainty in the literature.

Consistent with this interpretation, we find limited evidence for direct effects of environmental and management covariates. We tested the influence of a range of environmental and agronomic covariates. Contrary to previous studies suggesting stabilizing effects of inputs such as nitrogen [15], we find no consistent of fertilizers effect once mean–variance scaling is accounted for. This suggests that previously reported stabilizing effects may partly reflect impacts on mean yield rather than on variability itself. More broadly, the absence of significant covariate effects—other than broad crop categories—likely reflects both the limited size of the dataset and its bias toward temperate regions.

Beyond these general patterns, crop type emerges as an important source of variation, including when the effect of higher average yields values in these systems is taken into account. Crop type also plays an important role. Horticultural crops exhibit smaller and more uncertain yield gaps compared to cereals, and similar stability–productivity relationships between systems. This suggests relatively limited trade-offs for these crops and highlights their potential role in transitions toward more sustainable systems.

These results also offer a nuanced perspective on the performance of organic systems. Our results can be interpreted in contrasting ways for organic agriculture. On the one hand, it is notable that, despite lower investment in research and development and the reliance on varieties bred for conventional systems, organic systems exhibit a productivity–stability relationship comparable to that of conventional systems. On the other hand, the limited divergence in scaling relationships suggests that practices promoting soil quality or biodiversity do not translate into strong buffering effects on yield variability in the experimental settings analyzed here. This contrasts with some expectations from ecological theory and highlights the need for more empirical evidence at larger spatial and temporal scales.

Overall, our findings suggest that future research should focus on jointly analyzing productivity and variability across spatial scales to better inform the design of resilient agricultural systems.

## Supplementary Information

This Supplementary Information provides additional details on data compilation, variable definitions, statistical analyses, model comparison, diagnostic checks, and supplementary figures and tables supporting the main text.

## S1 Supplementary Methods

### S1.1 Data compilation

The dataset used in this study was compiled from a global database of experimental yield comparisons between organic, diversified and conventional agricultural systems [32]. This database was constructed through a two-step procedure combining (i) the merging of five previously published datasets and (ii) a systematic literature search conducted in Web of Science and Scopus. The initial compilation integrated data from major meta-analyses on organic and diversified farming systems, after removing duplicate records and studies lacking appropriate conventional controls. This was complemented by an updated systematic search using predefined Boolean queries targeting yield comparisons across farming systems. Articles were screened sequentially based on title, abstract and full text, and included only if they met strict criteria: (i) direct comparison between alternative (organic or diversified) and conventional systems, (ii) same crop species, site and year, and (iii) a minimum experimental duration of four years. This procedure resulted in a final dataset comprising 102 peer-reviewed studies. Data extraction included annual yield values and associated metadata (crop species, experimental design, management practices, and site characteristics). When necessary, data were digitized from figures or obtained directly from authors. Multiple quality-control steps were implemented throughout the process, including independent verification and consistency checks, ensuring the robustness and reproducibility of the database.

### S1.2 Dataset

The final dataset consists of 322 experimental units (EU), defined as the smallest scale of comparison between alternative and conventional systems for a given crop, site and year [32]. Each experimental unit represents a unique yield comparison under comparable agronomic conditions, thereby minimizing confounding effects. The dataset spans 73 experimental sites located across 23 countries and six continents, with a strong representation of Europe and North America. In total, 39 crop species are included and grouped into nine crop types, with cereals accounting for more than half of the observations. For each experimental unit, annual yield time series were compiled together with harmonized metadata describing agricultural practices (e.g., fertilization, irrigation, tillage, diversification strategies) and experimental context. From these data, standardized effect sizes were computed, including the log response ratio of mean yield, variance, and coefficient of variation, each associated with a measure of uncertainty. The structure of the dataset enables consistent comparisons of both yield levels and yield variability across a wide range of crops and management systems, and is specifically designed to support reproducible meta-analyses and the integration of future long-term experimental results.

### S1.3 Variables and derived metrics

We use the natural logarithm of the ratio of organic to conventional yields as an effect size metric. The use of ratios allows comparison across studies reporting yields in different units. The log response ratio is defined as:

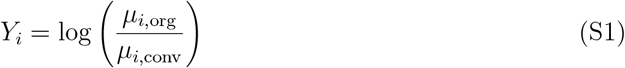

where *µ*_*i*,org_ and *µ*_*i*,conv_ are the mean yields in organic and conventional systems, respectively, computed across years within each experimental unit *i*.

To account for differences in precision between experimental units, we compute weights based on the variance of the log yield ratio. Following Nakagawa et al., this variance is:

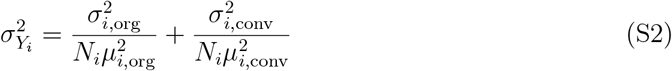

Where 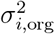 and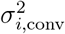are the interannual yield variances in organic and conventional systems,respectively, and *N*_*i*_ is the number of years in experimental unit *i*. In our dataset, *N*_*i*,org_ = *N*_*i*,conv_ in most cases, as yield time series are matched within experimental units (with *N*_*i*,org_ = *N*_*i*,conv_ = *N*_*i*_).

We then compute the corrected logarithm of the variance ratio:

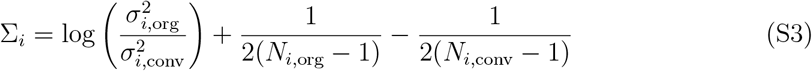

The variance of Σ_*i*_ is given by:

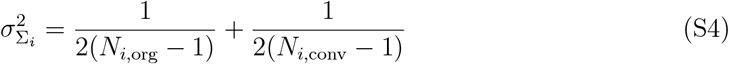

We also compute corrected logarithms of standard deviations:

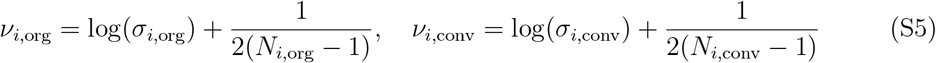

Finally, we consider the corrected logarithm of the coefficient of variation ratio:

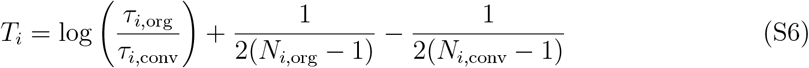

where

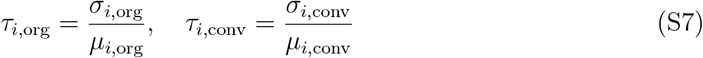

The variance of *T*_*i*_ is given by:

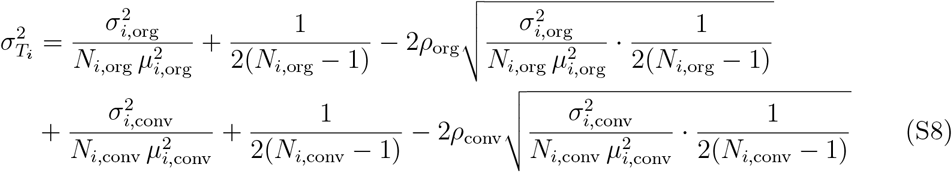

where *ρ*_org_ and *ρ*_conv_ are the correlations between *v*_*i*_ and *µ*_*i*_ in organic and conventional systems, respectively.

To summarize, we compute three main effect sizes: *Y*_*i*_ and its variance 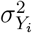 (yield ratio),Σ_*i*_ and its variance 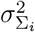 (variance ratio), and *T*_*i*_ and its variance 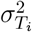 (coefficient of variation ratio).

### S1.4 Statistical modelling

#### Model comparisons

We compared a set of candidate MCMCglmm models to evaluate how yield variability (`logSdAll`) was related to management system (organic versus conventional; Group), mean yield (`logMeanAll`), and crop-type descriptors (`Types or CropTypes`). All models were fitted on the complete dataset and compared using the Deviance Information Criterion (DIC), where lower values indicate better model support. Models including experimental-unit random effects (`NbUnitAll`) and measurement-error weights consistently outperformed models using study-level random effects (`StudyAll`) or no weighting. The top-ranked models showed very similar support (ΔDIC < 2), indicating no clear single best model. The model including an interaction between mean yield and crop type (Group + `logMeanAll * Types;` M4) had the lowest DIC (599.41), closely followed by models including an interaction between cropping system and mean yield with crop-type covariates (M3 and M2; ΔDIC < 0.3). Given this model selection uncertainty, we retained model M3 (`Group* logMeanAll + CropTypes`) for inference because it provided a comparable fit to the data while including a larger number of statistically significant terms and accounting for crop-type variability using a more detailed classification. Model selection was therefore guided not only by DIC but also by interpretability and statistical support of individual terms.

Models without interactions, without weighting, or using study-level random effects showed substantially poorer fit (higher DIC values). Overall, these comparisons indicate that accounting for crop type improves model fit and that using experimental-unit random effects with weighting provides the most appropriate description of variation in yield variability across systems.

Weakly informative priors were specified for all models using a standard formulation in `MCMCglmm`. Fixed effects were assigned diffuse Gaussian priors with mean zero and a large variance (*V* = 10^6^), reflecting minimal prior information. For the random effects and residual variance components, inverse-Wishart priors were used with scale parameter *V* = 1 and low degrees of freedom (*v* = 1), corresponding to weakly informative priors with broad support. This choice allows the data to dominate posterior estimation while ensuring model stability. Sensitivity analyses using alternative values of *v* (e.g., *v* = 0.002) yielded qualitatively similar results, indicating that inference was robust to prior specification.

**Table S1:**
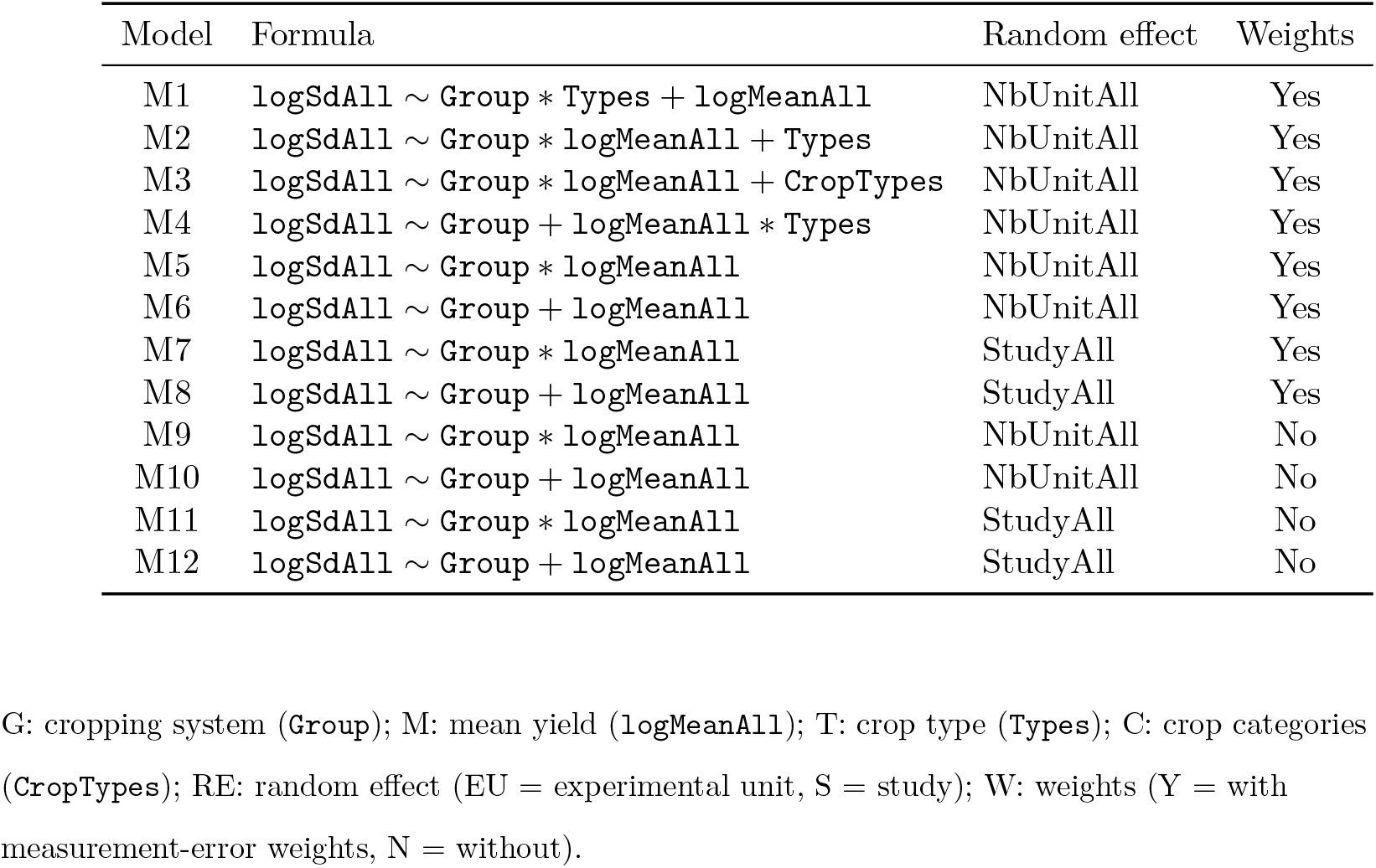
Definition of candidate models.

**Table S2:**
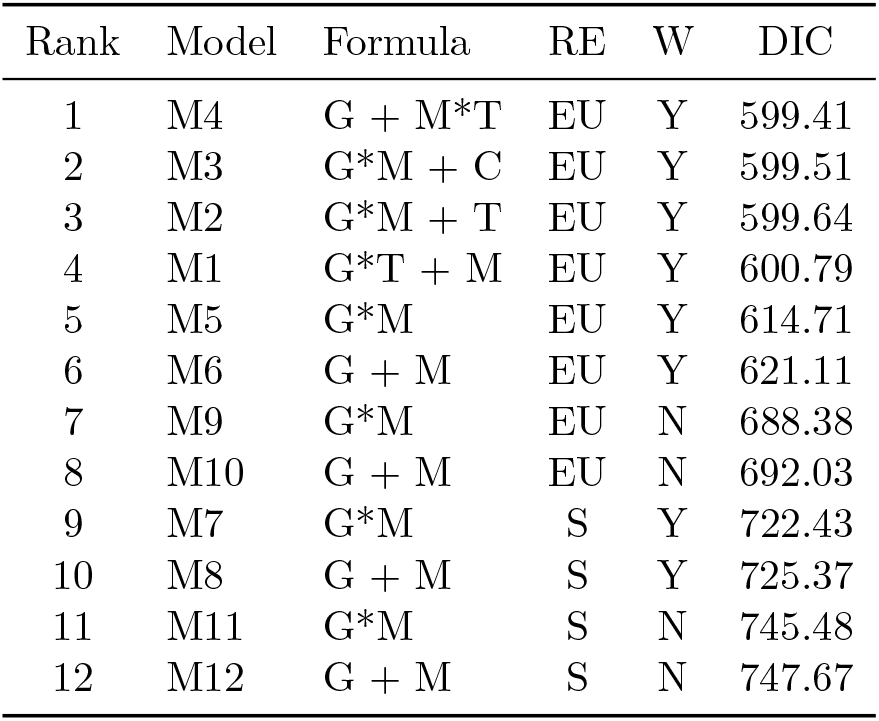
Comparison of candidate models ranked by increasing DIC.

The top-ranked models showed very similar support (ΔDIC < 2), indicating no clear single best model. The model including an interaction between mean yield and crop type (G + M*×*T) had the lowest DIC, but models including interactions between cropping system and mean yield with crop-type covariates also received comparable support.

#### Convergence diagnostics for M3

Basic convergence diagnostics were examined for the variance components of the selected MCMCglmm model using trace plots, posterior density plots, autocorrelation diagnostics, Heidelberger–Welch diagnostics, and effective sample sizes. Visual inspection of the chains indicated satisfactory mixing and no obvious long-term drift for the variance components associated with the experimental unit and the residual term. Their posterior densities were unimodal and approximately symmetric, supporting stable posterior estimation. By contrast, the measurement-error component entered the model through the mev argument as a fixed variance structure rather than a free parameter to be estimated; accordingly, it remained effectively constant across iterations and was not interpreted as part of the convergence assessment. Overall, the diagnostics support satisfactory convergence of the estimated variance components in the selected model.

### S1.5 Sensitivity analysis

Sensitivity analyses were conducted by varying prior specifications for the variance components, notably using lower degrees of freedom in the inverse-Wishart distributions (e.g., *v* = 0.002). These alternative specifications yielded qualitatively similar parameter estimates and model rankings, indicating that the main conclusions were robust to prior choice.

## S2 Supplementary Equations

### S2.1 Effect size definitions

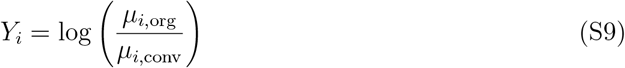

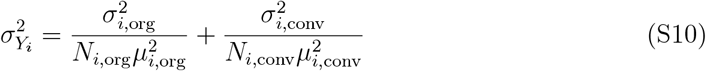

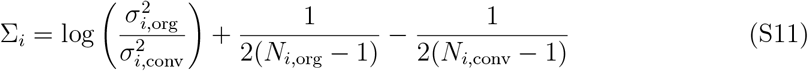

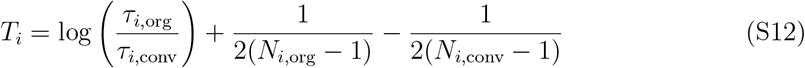

### S2.2 Taylor power law

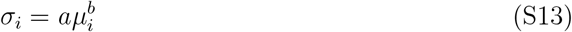

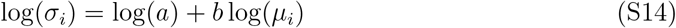

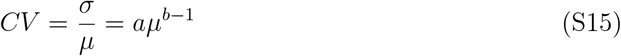

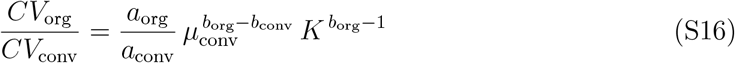

where

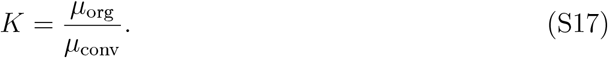

## S3 Supplementary Results

### S3.1 Overall yield ratio between organic and conventional systems

We first estimate the overall difference in mean yields between organic/diversified and conventional systems using an intercept-only Bayesian mixed-effects model, with study number included as a random effect to account for non-independence among observations within studies. Organic and diversified are not separated because of the relatively small number of data points for diversified systems. The model provides an estimate of the average log response ratio across all experimental units. The posterior mean of the log yield ratio is negative and significantly different from zero (posterior mean: −0.158, 95% credible interval: [−0.223, −0.093], *p <* 0.001), indicating that organic systems produce lower yields on average than conventional systems. Back-transformation of this estimate yields an average organic-to-conventional yield ratio of approximately 0.85 [0.80, 0.91], corresponding to a mean yield gap of 14.6% [8.9%, 20.0%]. These results confirm the presence of a systematic yield gap in the dataset.

**Table S3:**
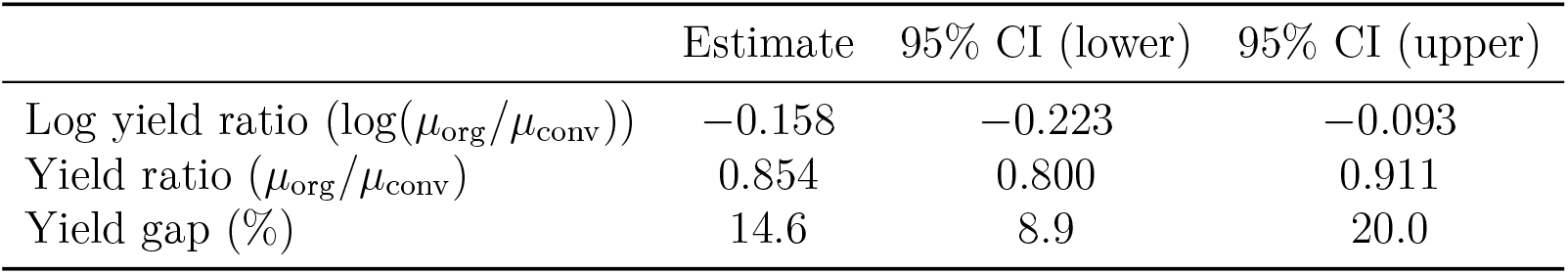
Summary of the estimated organic-to-conventional yield ratio from the intercept-only model.

### S3.2 Overall variance ratio between organic and conventional systems

We next estimate the overall difference in yield variability between organic and conventional systems using an intercept-only Bayesian mixed-effects model, with study identity included as a random effect. The model provides an estimate of the average log variance ratio across all experimental units. The posterior mean of the log variance ratio is close to zero and not significantly different from zero (posterior mean: 0.007, [−0.106, 0.112], *p* = 0.91), indicating no systematic difference in yield variability between organic and conventional systems. Back-transformation yields a variance ratio of approximately 1.01 [0.90, 1.12], suggesting that variability levels are, on average, comparable across systems. These results indicate that, despite the presence of a clear yield gap, absolute variability does not differ significantly between management systems when considered independently of mean yield.

**Table S4:**
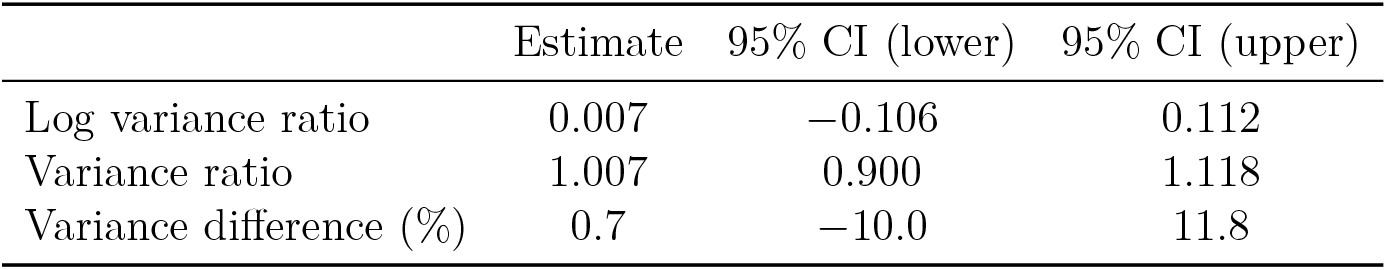
Summary of the estimated variance ratio between organic and conventional systems.

### S3.3 Overall coefficient of variation ratio between systems

We finally estimate differences in relative yield variability using the coefficient of variation (CV), again using an intercept-only Bayesian mixed-effects model with study identity as a random effect. This model provides an estimate of the average log CV ratio across all experimental units. In contrast to the variance ratio, the log CV ratio is positive and significantly different from zero (posterior mean: 0.168 [0.064, 0.267], *p* = 0.002), indicating that organic systems exhibit higher relative variability than conventional systems. Back-transformation yields a CV ratio of approximately 1.18 [1.07, 1.31], corresponding to an average increase of about 18% in relative variability under organic management. Taken at face value, this result would suggest that organic systems are less stable than conventional systems. However, when interpreted jointly with the absence of differences in absolute variability and the presence of a substantial yield gap, this pattern is consistent with a mechanical effect arising from the dependence of the CV on mean yield. As such, the higher CV observed in organic systems does not necessarily reflect greater intrinsic variability, but rather results from lower mean yields combined with comparable levels of variance.

**Table S5:**
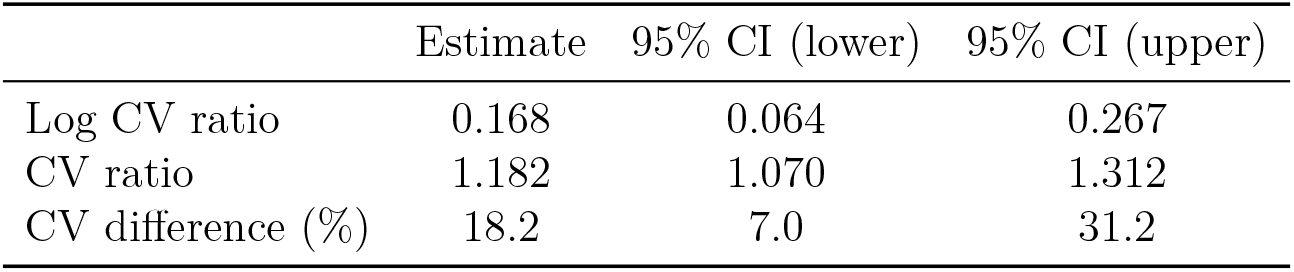
Summary of the estimated coefficient of variation ratio between organic and conventional systems.

## S4 Supplementary Figures

Forest plots of yield, variance, and CV ratios for grain and garden crops are provided below.

**Figure S1:**
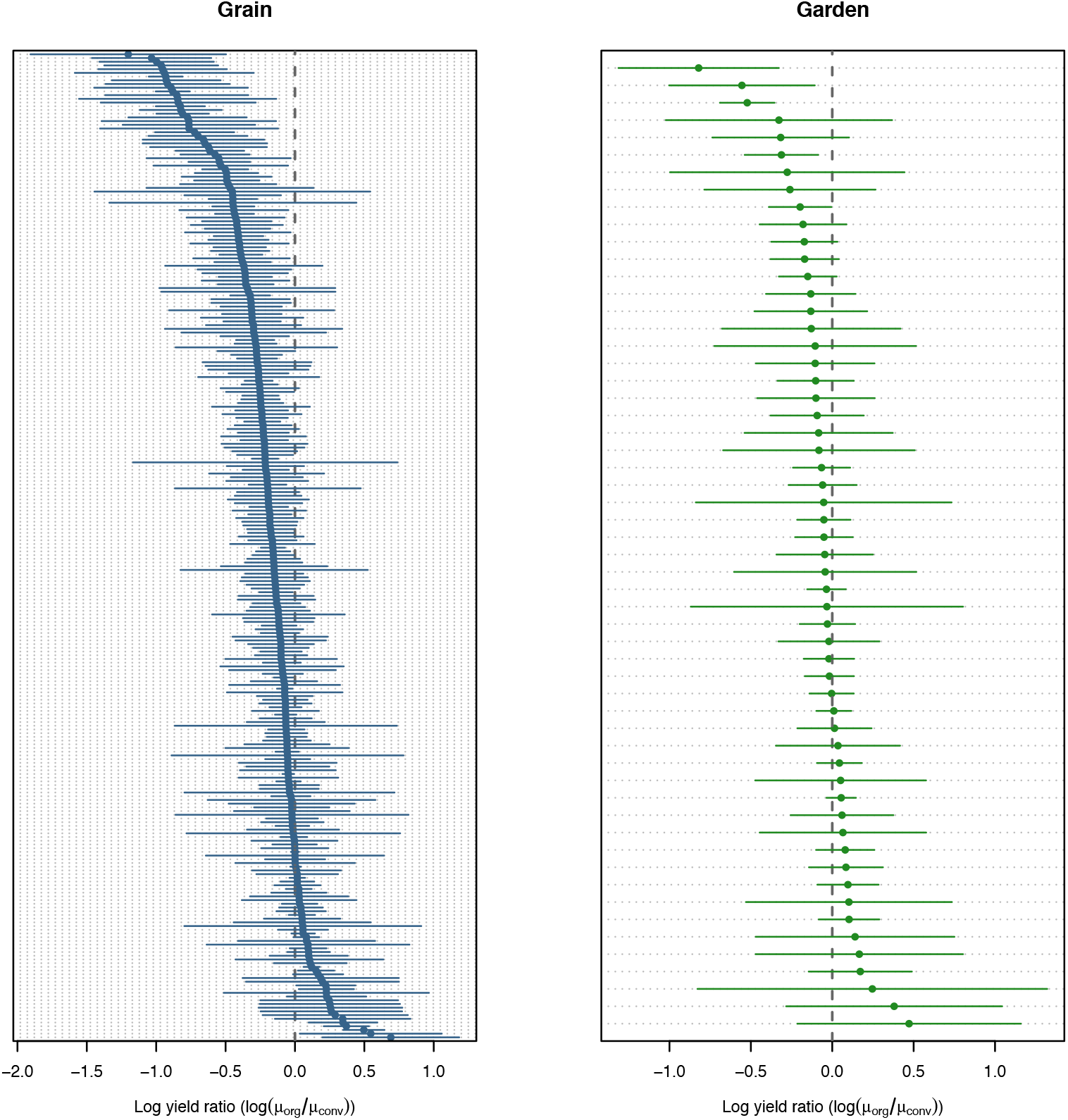
Forest plot of the organic-to-conventional yield log ratio for grain (left) and garden (right) crops independently. Each point corresponds to a single experimental unit (EU). Confidence intervals are computed based on the equations presented in Section S2. The gray dotted line indicates equality between organic and conventional yields.

**Figure S2:**
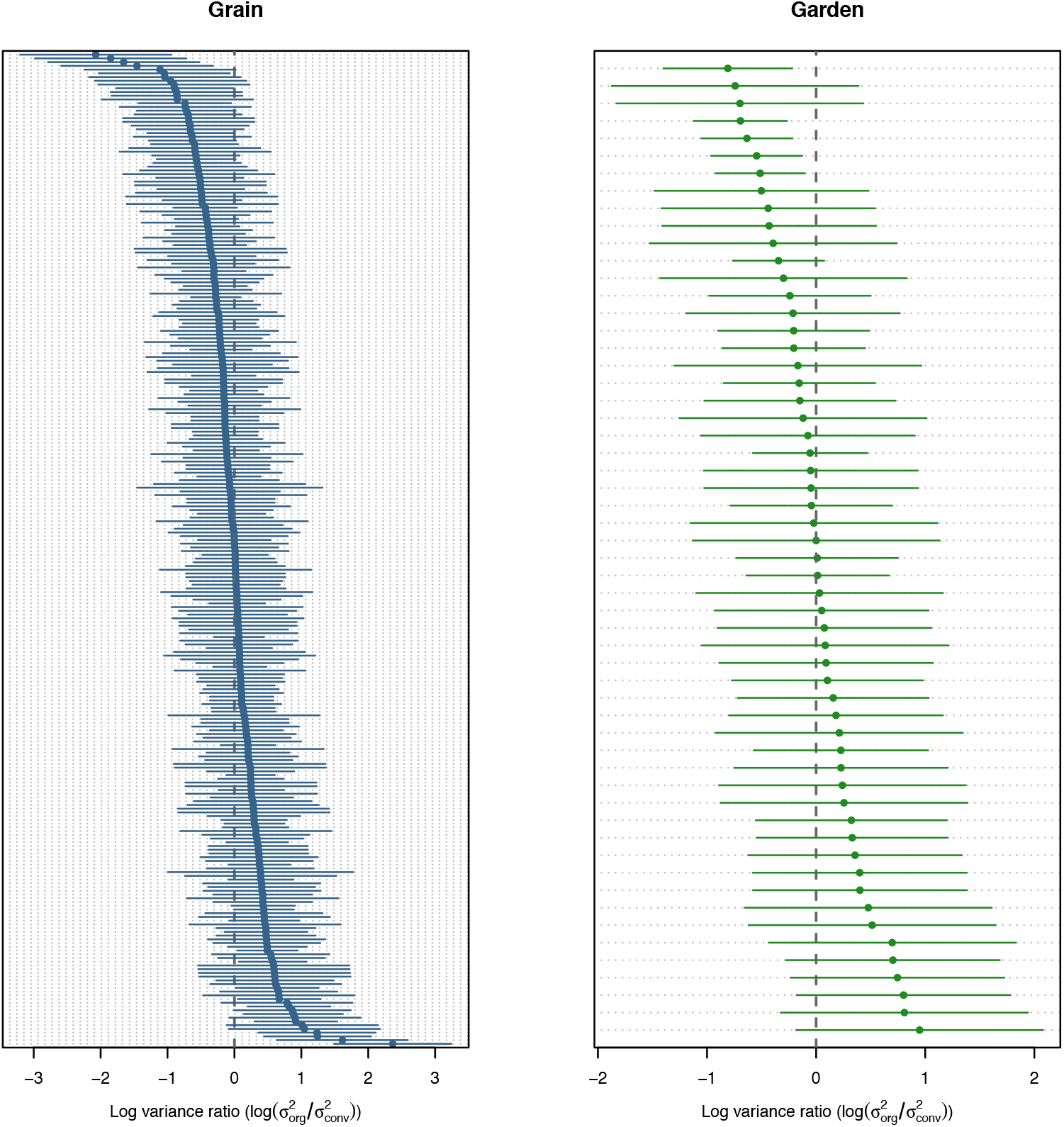
Forest plot of the organic-to-conventional variance log ratio for grain (left) and garden (right) crops independently. Each point corresponds to a single experimental unit (EU). Confidence intervals are computed based on the equations presented in Section S2. The gray dotted line indicates equality between organic and conventional yield variances.

**Figure S3:**
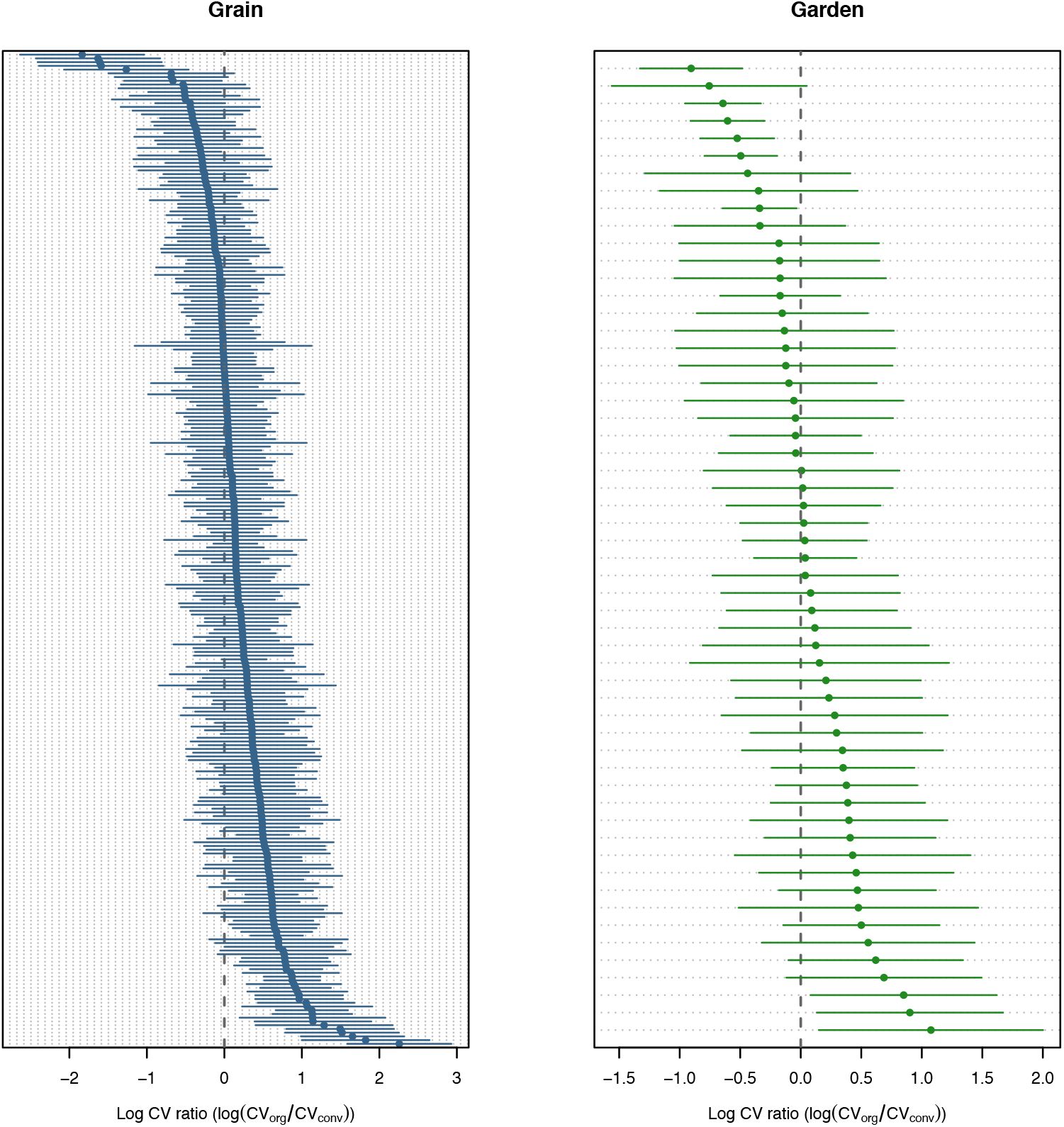
Forest plot of the organic-to-conventional CV log ratio for grain (left) and garden (right) crops independently. Each point corresponds to a single experimental unit (EU). Confidence intervals are computed based on the equations presented in Section S2. The gray dotted line indicates equality between organic and conventional yield CV.

**Figure S4:**
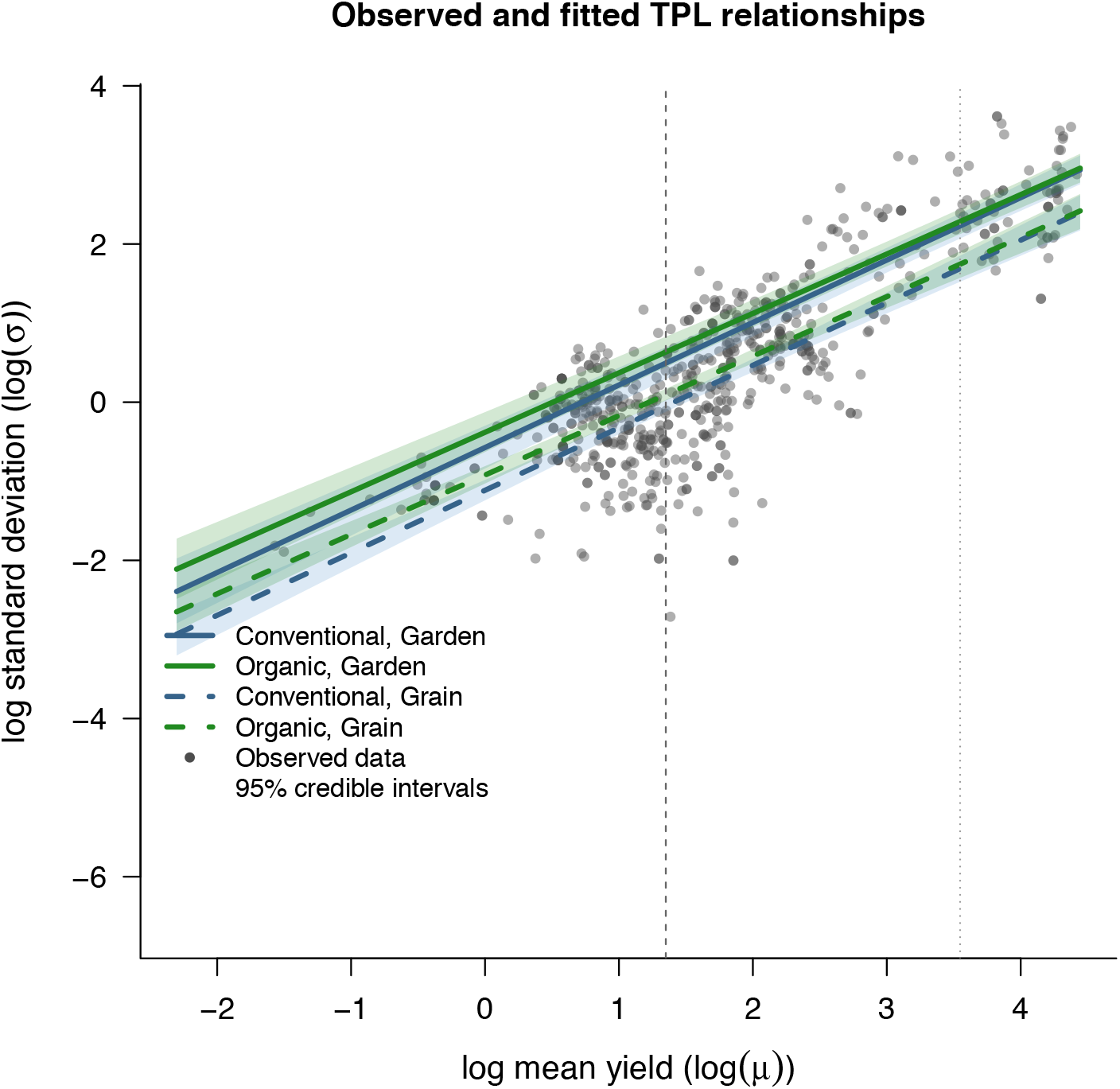
Taylor Power Law as estimated from the best statistical model (see Figure 4 in organic and conventional systems for grain and garden systems. Shaded areas indicate 95% credible intervals for each system independently.

**Figure S5:**
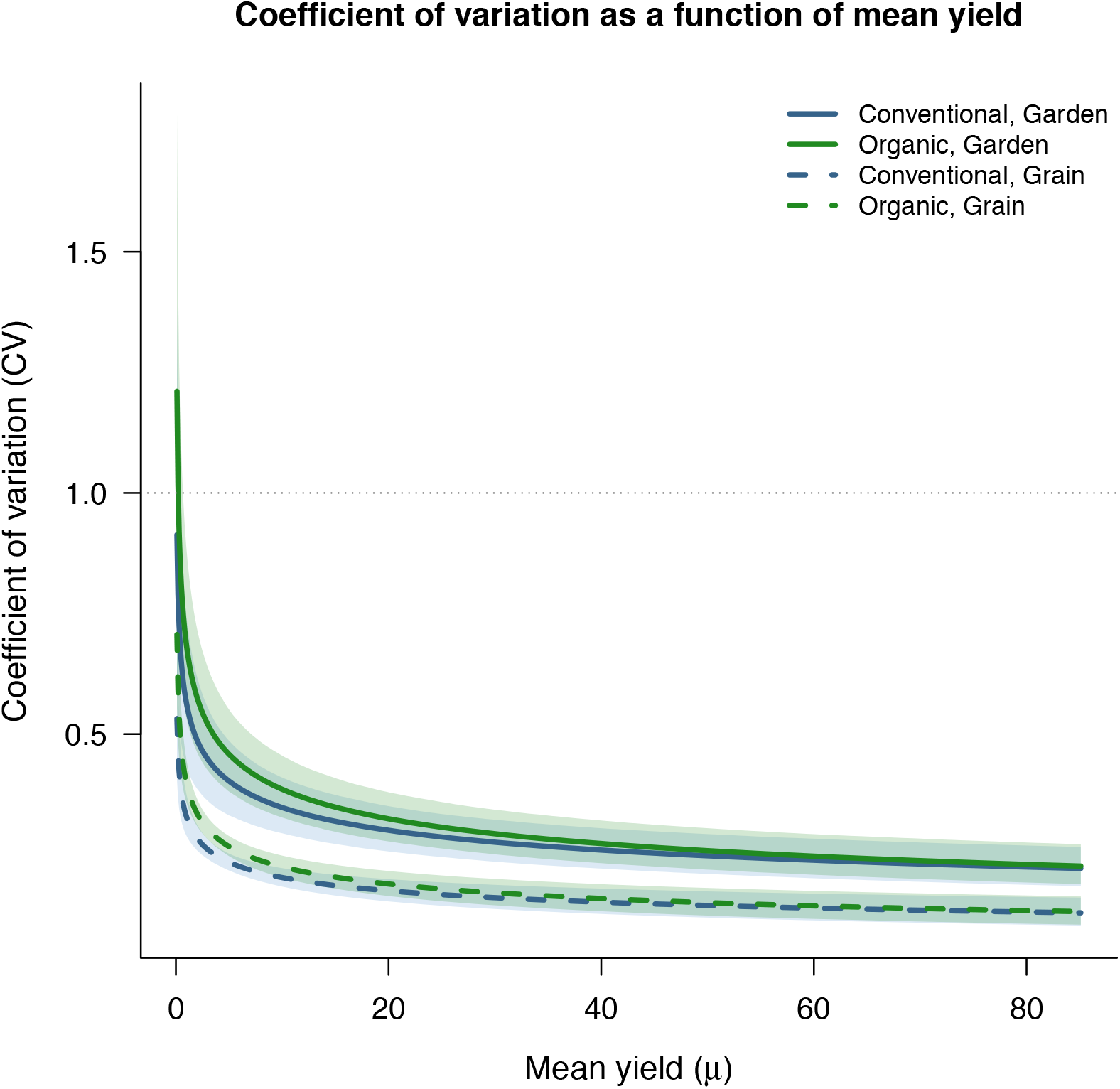
Relationship between the coefficient of variation (CV) and mean yields (t/ha) in organic and conventional systems for grain and garden systems. Shaded areas indicate 95% credible intervals. The horizontal gray lines at 1 delineate conditions where organic systems have lower or higher yields and variability than conventional systems. In both crop groups, the CV ratio decreases as the yield ratio increases, indicating that apparent differences in stability are strongly driven by differences in mean yield.

