## Supplementary material and Figures for "Mean yield gap explains differences in yield stability between organic and conventional systems"

$$Y_i = \log\left(\frac{\mu_{i,\mathrm{org}}}{\mu_{i,\mathrm{conv}}}\right) \tag{S1}$$

where $\mu_{i,\mathrm{org}}$ and $\mu_{i,\mathrm{conv}}$ are the mean yields in organic and conventional systems, respectively, computed across years within each experimental unit $i$.

To account for differences in precision between experimental units, we compute weights based on the variance of the log yield ratio. Following Nakagawa et al., this variance is:

$$\sigma^2_{Y_i} = \frac{\sigma^2_{i,\mathrm{org}}}{N_i \mu^2_{i,\mathrm{org}}} + \frac{\sigma^2_{i,\mathrm{conv}}}{N_i \mu^2_{i,\mathrm{conv}}} \tag{S2}$$

where $\sigma^2_{i,\mathrm{org}}$ and $\sigma^2_{i,\mathrm{conv}}$ are the interannual yield variances in organic and conventional systems, respectively, and $N_i$ is the number of years in experimental unit $i$. In our dataset, $N_{i,\mathrm{org}} = N_{i,\mathrm{conv}}$ in most cases, as yield time series are matched within experimental units (with $N_{i,\mathrm{org}} = N_{i,\mathrm{conv}} = N_i$).

We then compute the corrected logarithm of the variance ratio:

$$\Sigma_i = \log\left(\frac{\sigma^2_{i,\mathrm{org}}}{\sigma^2_{i,\mathrm{conv}}}\right) + \frac{1}{2(N_{i,\mathrm{org}} - 1)} - \frac{1}{2(N_{i,\mathrm{conv}} - 1)} \tag{S3}$$

The variance of $\Sigma_i$ is given by:

$$\sigma^2_{\Sigma_i} = \frac{1}{2(N_{i,\mathrm{org}} - 1)} + \frac{1}{2(N_{i,\mathrm{conv}} - 1)} \tag{S4}$$

We also compute corrected logarithms of standard deviations:

$$\nu_{i,\mathrm{org}} = \log(\sigma_{i,\mathrm{org}}) + \frac{1}{2(N_{i,\mathrm{org}} - 1)}, \quad \nu_{i,\mathrm{conv}} = \log(\sigma_{i,\mathrm{conv}}) + \frac{1}{2(N_{i,\mathrm{conv}} - 1)} \tag{S5}$$

Finally, we consider the corrected logarithm of the coefficient of variation ratio:

$$T_i = \log\left(\frac{\tau_{i,\mathrm{org}}}{\tau_{i,\mathrm{conv}}}\right) + \frac{1}{2(N_{i,\mathrm{org}} - 1)} - \frac{1}{2(N_{i,\mathrm{conv}} - 1)} \tag{S6}$$

where

$$\tau_{i,\mathrm{org}} = \frac{\sigma_{i,\mathrm{org}}}{\mu_{i,\mathrm{org}}}, \quad \tau_{i,\mathrm{conv}} = \frac{\sigma_{i,\mathrm{conv}}}{\mu_{i,\mathrm{conv}}} \tag{S7}$$

The variance of $T_i$ is given by:

$$\sigma^2_{T_i} = \frac{\sigma^2_{i,\text{org}}}{N_{i,\text{org}}\,\mu^2_{i,\text{org}}} + \frac{1}{2(N_{i,\text{org}}-1)} - 2\rho_{\text{org}}\sqrt{\frac{\sigma^2_{i,\text{org}}}{N_{i,\text{org}}\,\mu^2_{i,\text{org}}}\cdot\frac{1}{2(N_{i,\text{org}}-1)}} + \frac{\sigma^2_{i,\text{conv}}}{N_{i,\text{conv}}\,\mu^2_{i,\text{conv}}} + \frac{1}{2(N_{i,\text{conv}}-1)} - 2\rho_{\text{conv}}\sqrt{\frac{\sigma^2_{i,\text{conv}}}{N_{i,\text{conv}}\,\mu^2_{i,\text{conv}}}\cdot\frac{1}{2(N_{i,\text{conv}}-1)}} \quad \text{(S8)}$$

where $\rho_{\text{org}}$ and $\rho_{\text{conv}}$ are the correlations between $\nu_i$ and $\mu_i$ in organic and conventional systems, respectively.

To summarize, we compute three main effect sizes: $Y_i$ and its variance $\sigma^2_{Y_i}$ (yield ratio), $\Sigma_i$ and its variance $\sigma^2_{\Sigma_i}$ (variance ratio), and $T_i$ and its variance $\sigma^2_{T_i}$ (coefficient of variation ratio).

Table S1: Definition of candidate models.

| Model | Formula | Random effect | Weights |
|---|---|---|---|
| M1 | logSdAll ~ Group * Types + logMeanAll | NbUnitAll | Yes |
| M2 | logSdAll ~ Group * logMeanAll + Types | NbUnitAll | Yes |
| M3 | logSdAll ~ Group * logMeanAll + CropTypes | NbUnitAll | Yes |
| M4 | logSdAll ~ Group + logMeanAll * Types | NbUnitAll | Yes |
| M5 | logSdAll ~ Group * logMeanAll | NbUnitAll | Yes |
| M6 | logSdAll ~ Group + logMeanAll | NbUnitAll | Yes |
| M7 | logSdAll ~ Group * logMeanAll | StudyAll | Yes |
| M8 | logSdAll ~ Group + logMeanAll | StudyAll | Yes |
| M9 | logSdAll ~ Group * logMeanAll | NbUnitAll | No |
| M10 | logSdAll ~ Group + logMeanAll | NbUnitAll | No |
| M11 | logSdAll ~ Group * logMeanAll | StudyAll | No |
| M12 | logSdAll ~ Group + logMeanAll | StudyAll | No |

G: cropping system (`Group`); M: mean yield (`logMeanAll`); T: crop type (`Types`); C: crop categories (`CropTypes`); RE: random effect (EU = experimental unit, S = study); W: weights (Y = with measurement-error weights, N = without).

Table S2: Comparison of candidate models ranked by increasing DIC.

| Rank | Model | Formula | RE | W | DIC |
|---|---|---|---|---|---|
| 1 | M4 | G + M*T | EU | Y | 599.41 |
| 2 | M3 | G*M + C | EU | Y | 599.51 |
| 3 | M2 | G*M + T | EU | Y | 599.64 |
| 4 | M1 | G*T + M | EU | Y | 600.79 |
| 5 | M5 | G*M | EU | Y | 614.71 |
| 6 | M6 | G + M | EU | Y | 621.11 |
| 7 | M9 | G*M | EU | N | 688.38 |
| 8 | M10 | G + M | EU | N | 692.03 |
| 9 | M7 | G*M | S | Y | 722.43 |
| 10 | M8 | G + M | S | Y | 725.37 |
| 11 | M11 | G*M | S | N | 745.48 |
| 12 | M12 | G + M | S | N | 747.67 |

The top-ranked models showed very similar support ($\Delta$DIC $< 2$), indicating no clear single best model. The model including an interaction between mean yield and crop type (G + M$\times$T) had the lowest DIC, but models including interactions between cropping system and mean yield with crop-type covariates also received comparable support.

**Convergence diagnostics for M3** Basic convergence diagnostics were examined for the variance components of the selected MCMCglmm model using trace plots, posterior density plots, autocorrelation diagnostics, Heidelberger–Welch diagnostics, and effective sample sizes. Visual inspection of the chains indicated satisfactory mixing and no obvious long-term drift for the variance components associated with the experimental unit and the residual term. Their posterior densities were unimodal and approximately symmetric, supporting stable posterior estimation. By contrast, the measurement-error component entered the model through the `mev` argument as a fixed variance structure rather than a free parameter to be estimated; accordingly, it remained effectively constant across iterations and was not interpreted as part of the convergence assessment. Overall, the diagnostics support satisfactory convergence of the estimated variance components in the selected model.

##### S1.5 Sensitivity analysis

Sensitivity analyses were conducted by varying prior specifications for the variance components, notably using lower degrees of freedom in the inverse-Wishart distributions (e.g., $\nu = 0.002$). These alternative specifications yielded qualitatively similar parameter estimates and model rankings, indicating that the main conclusions were robust to prior choice.

#### S2 Supplementary Equations

##### S2.1 Effect size definitions

$$Y_i = \log\left(\frac{\mu_{i,\mathrm{org}}}{\mu_{i,\mathrm{conv}}}\right) \tag{S9}$$

$$\sigma^2_{Y_i} = \frac{\sigma^2_{i,\mathrm{org}}}{N_{i,\mathrm{org}}\mu^2_{i,\mathrm{org}}} + \frac{\sigma^2_{i,\mathrm{conv}}}{N_{i,\mathrm{conv}}\mu^2_{i,\mathrm{conv}}} \tag{S10}$$

$$\Sigma_i = \log\left(\frac{\sigma^2_{i,\mathrm{org}}}{\sigma^2_{i,\mathrm{conv}}}\right) + \frac{1}{2(N_{i,\mathrm{org}} - 1)} - \frac{1}{2(N_{i,\mathrm{conv}} - 1)} \tag{S11}$$

$$T_i = \log\left(\frac{\tau_{i,\mathrm{org}}}{\tau_{i,\mathrm{conv}}}\right) + \frac{1}{2(N_{i,\mathrm{org}} - 1)} - \frac{1}{2(N_{i,\mathrm{conv}} - 1)} \tag{S12}$$

##### S2.2 Taylor power law

$$\sigma_i = a\mu_i^b \tag{S13}$$

$$\log(\sigma_i) = \log(a) + b\log(\mu_i) \tag{S14}$$

$$CV = \frac{\sigma}{\mu} = a\mu^{b-1} \tag{S15}$$

$$\frac{CV_{\text{org}}}{CV_{\text{conv}}} = \frac{a_{\text{org}}}{a_{\text{conv}}} \, \mu_{\text{conv}}^{b_{\text{org}} - b_{\text{conv}}} \, K^{\, b_{\text{org}} - 1} \tag{S16}$$

where

$$K = \frac{\mu_{\text{org}}}{\mu_{\text{conv}}}. \tag{S17}$$

### S3 Supplementary Results

#### S3.1 Overall yield ratio between organic and conventional systems

We first estimate the overall difference in mean yields between organic/diversified and conventional systems using an intercept-only Bayesian mixed-effects model, with study number included as a random effect to account for non-independence among observations within studies. Organic and diversified are not separated because of the relatively small number of data points for diversified systems. The model provides an estimate of the average log response ratio across all experimental units. The posterior mean of the log yield ratio is negative and significantly different from zero (posterior mean: −0.158, 95% credible interval: [−0.223, −0.093], $p < 0.001$), indicating that organic systems produce lower yields on average than conventional systems. Back-transformation of this estimate yields an average organic-to-conventional yield ratio of approximately 0.85 [0.80, 0.91], corresponding to a mean yield gap of 14.6% [8.9%, 20.0%]. These results confirm the presence of a systematic yield gap in the dataset.

Table S3: Summary of the estimated organic-to-conventional yield ratio from the intercept-only model.

| | Estimate | 95% CI (lower) | 95% CI (upper) |
|---|---|---|---|
| Log yield ratio ($\log(\mu_{\text{org}}/\mu_{\text{conv}})$) | −0.158 | −0.223 | −0.093 |
| Yield ratio ($\mu_{\text{org}}/\mu_{\text{conv}}$) | 0.854 | 0.800 | 0.911 |
| Yield gap (%) | 14.6 | 8.9 | 20.0 |

#### S3.2 Overall variance ratio between organic and conventional systems

We next estimate the overall difference in yield variability between organic and conventional systems using an intercept-only Bayesian mixed-effects model, with study identity included as a random effect. The model provides an estimate of the average log variance ratio across all experimental units. The posterior mean of the log variance ratio is close to zero and not significantly different from zero (posterior mean: 0.007, $[-0.106, 0.112]$, $p = 0.91$), indicating no systematic difference in yield variability between organic and conventional systems. Back-transformation yields a variance ratio of approximately 1.01 [0.90, 1.12], suggesting that variability levels are, on average, comparable across systems. These results indicate that, despite the presence of a clear yield gap, absolute variability does not differ significantly between management systems when considered independently of mean yield.

Table S4: Summary of the estimated variance ratio between organic and conventional systems.

| | Estimate | 95% CI (lower) | 95% CI (upper) |
|---|---|---|---|
| Log variance ratio | 0.007 | −0.106 | 0.112 |
| Variance ratio | 1.007 | 0.900 | 1.118 |
| Variance difference (%) | 0.7 | −10.0 | 11.8 |

#### S3.3 Overall coefficient of variation ratio between systems

We finally estimate differences in relative yield variability using the coefficient of variation (CV), again using an intercept-only Bayesian mixed-effects model with study identity as a random effect. This model provides an estimate of the average log CV ratio across all experimental units. In contrast to the variance ratio, the log CV ratio is positive and significantly different from zero (posterior mean: 0.168 [0.064, 0.267], $p = 0.002$), indicating that organic systems exhibit higher relative variability than conventional systems. Back-transformation yields a CV ratio of approximately 1.18 [1.07, 1.31], corresponding to an average increase of about 18% in relative variability under organic management. Taken at face value, this result would suggest that organic systems are less stable than conventional systems. However, when interpreted jointly with the absence of differences in absolute variability and the presence of a substantial yield gap, this pattern is consistent with a mechanical effect arising from the dependence of the CV on mean yield. As such, the higher CV observed in organic systems does not necessarily reflect greater intrinsic variability, but rather results from lower mean yields combined with comparable levels of variance.

Table S5: Summary of the estimated coefficient of variation ratio between organic and conventional systems.

| | Estimate | 95% CI (lower) | 95% CI (upper) |
|---|---|---|---|
| Log CV ratio | 0.168 | 0.064 | 0.267 |
| CV ratio | 1.182 | 1.070 | 1.312 |
| CV difference (%) | 18.2 | 7.0 | 31.2 |

#### S4 Supplementary Figures

Forest plots of yield, variance, and CV ratios for grain and garden crops are provided below.

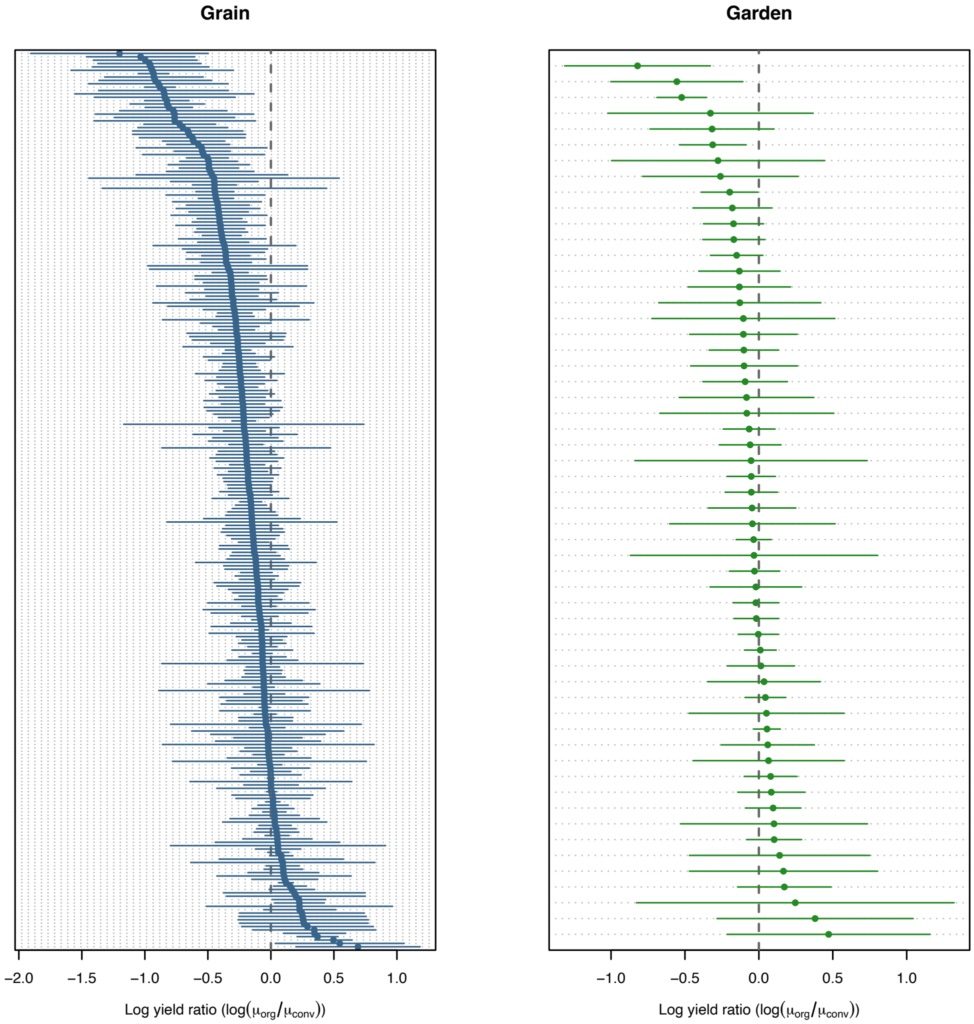


Figure S1: Forest plot of the organic-to-conventional yield log ratio for grain (left) and garden (right) crops independently. Each point corresponds to a single experimental unit (EU). Confidence intervals are computed based on the equations presented in Section S2. The gray dotted line indicates equality between organic and conventional yields.

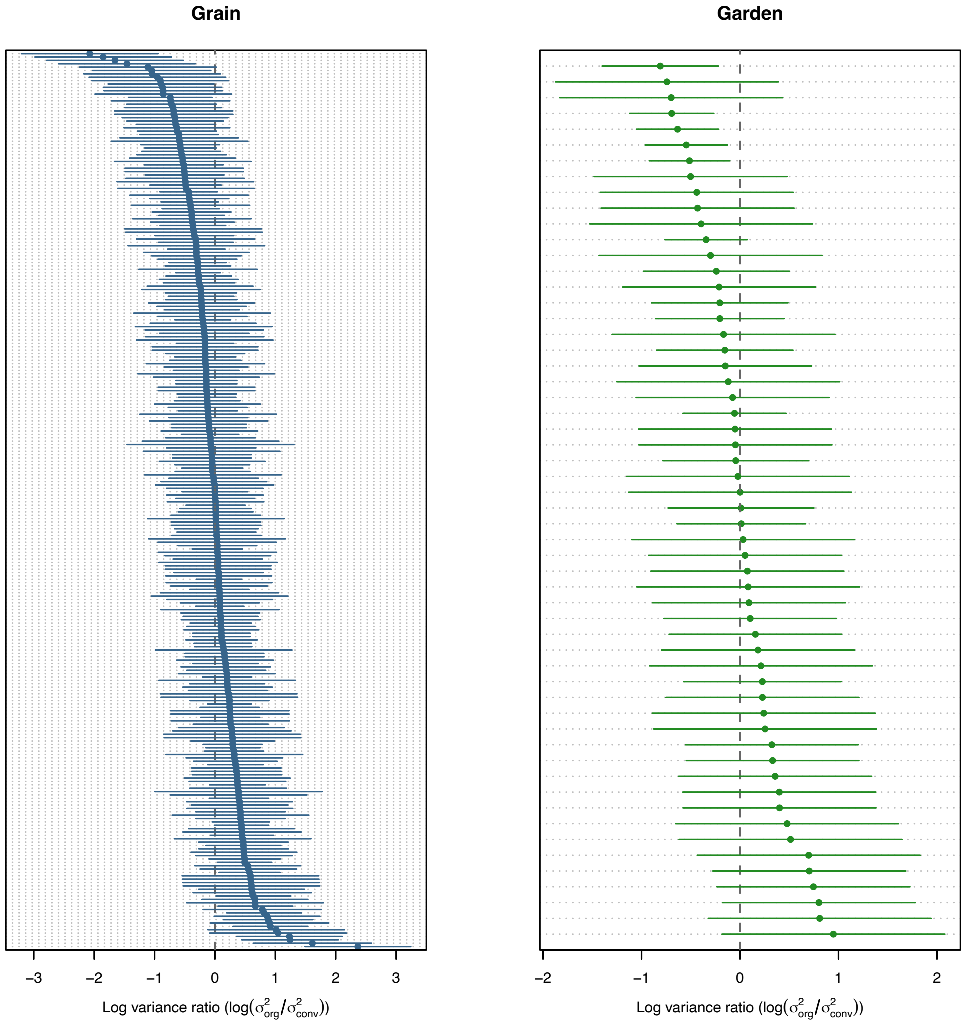


Figure S2: Forest plot of the organic-to-conventional variance log ratio for grain (left) and garden (right) crops independently. Each point corresponds to a single experimental unit (EU). Confidence intervals are computed based on the equations presented in Section S2. The gray dotted line indicates equality between organic and conventional yield variances.

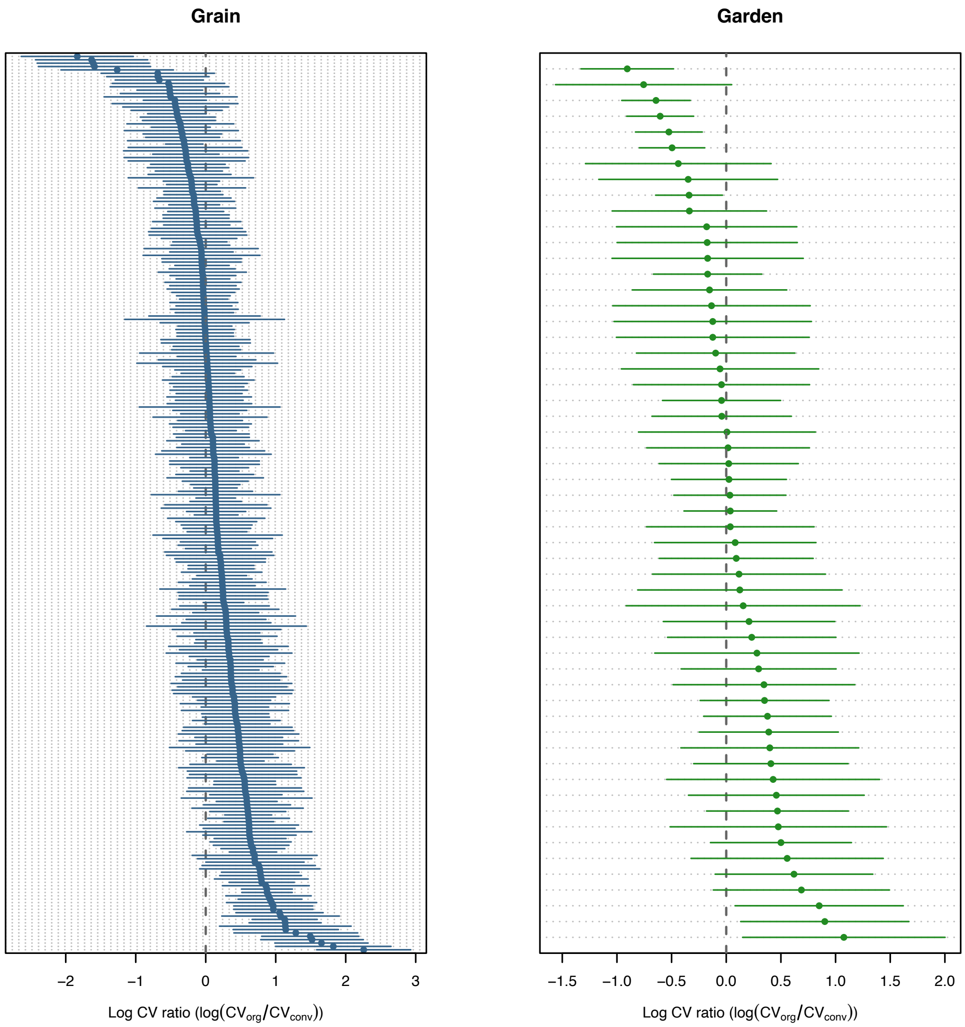


Figure S3: Forest plot of the organic-to-conventional CV log ratio for grain (left) and garden (right) crops independently. Each point corresponds to a single experimental unit (EU). Confidence intervals are computed based on the equations presented in Section S2. The gray dotted line indicates equality between organic and conventional yield CV.

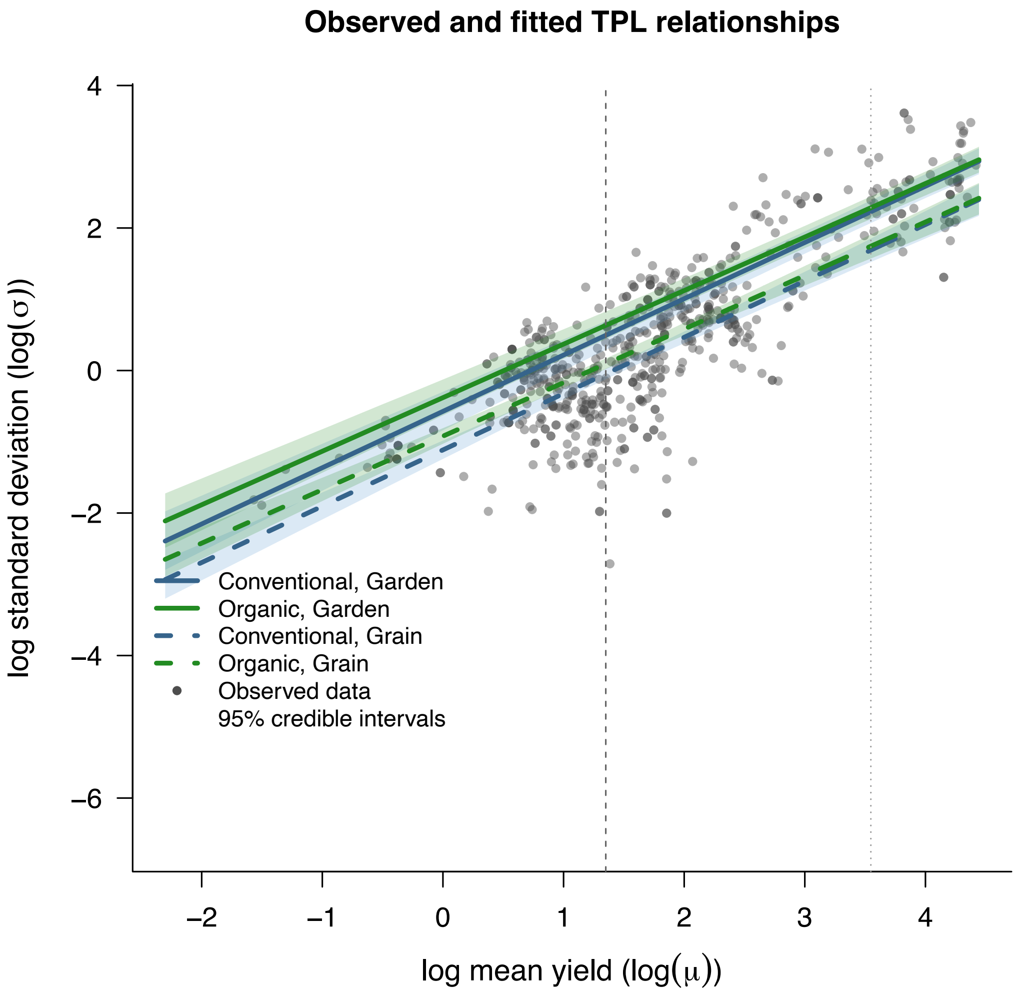


Figure S4: Taylor Power Law as estimated from the best statistical model (see Figure 4 in organic and conventional systems for grain and garden systems. Shaded areas indicate 95% credible intervals for each system independently.

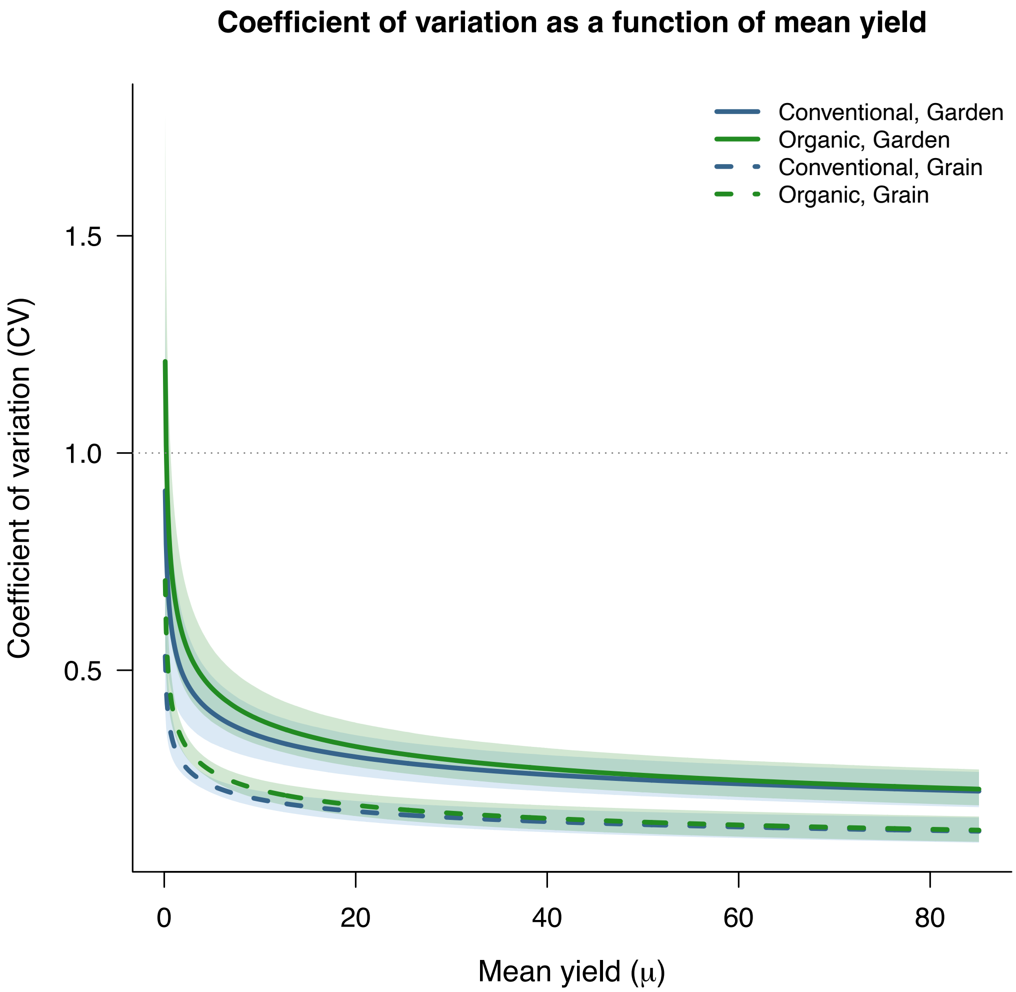


Figure S5: Relationship between the coefficient of variation (CV) and mean yields (t/ha) in organic and conventional systems for grain and garden systems. Shaded areas indicate 95% credible intervals. The horizontal gray lines at 1 delineate conditions where organic systems have lower or higher yields and variability than conventional systems. In both crop groups, the CV ratio decreases as the yield ratio increases, indicating that apparent differences in stability are strongly driven by differences in mean yield.
